# Disentangling the role of NAc D1 and D2 cells in hedonic eating

**DOI:** 10.1101/2022.10.29.514339

**Authors:** Mathilde C. C. Guillaumin, Paulius Viskaitis, Ed Bracey, Denis Burdakov, Daria Peleg-Raibstein

## Abstract

Overeating is driven by both the hedonic component (‘liking’) of food, and the motivation (‘wanting’) to eat it. The nucleus accumbens (NAc) is a key brain center implicated in these processes, but how distinct NAc cell populations encode ‘liking’ and ‘wanting’ to shape overconsumption remains unclear. Here, we probed the roles of NAc D1 and D2 cells in these processes using cell-specific recording and control in diverse behavioral paradigms that disentangle reward traits of ‘liking’ and ‘wanting’ related to food choice and overeating. NAc D2 cells encoded experience-dependent development of ‘liking’, while NAc D1 cells encoded innate ‘liking’ during the first food taste. Optogenetic control confirmed causal links of D1 and D2 cells to these aspects of ‘liking’. In relation to ‘wanting’, D1 and D2 cells encoded and promoted distinct aspects of food approach: D1 cells interpreted food cues while D2 cells also sustained food-visit-length that facilitates consumption. Finally, at the level of food choice, D1, but not D2, cell activity was sufficient to switch food preference, programming subsequent long-lasting overconsumption. By revealing complementary roles of D1 and D2 cells in consumption, these findings assign neural bases to ‘liking’ and ‘wanting’ in a unifying framework of D1 and D2 cell activity.

## Introduction

Overweight and obesity, and eating disorders generally, have increased in recent decades [1, 2]. Thus, it is crucial to better understand the factors that contribute to the development of different eating disorders. In the modern obesogenic environment, palatability and availability of foods play a major role in the development of overeating disorders [3, 4]. Food consumption is not only motivated by the homeostatic need to monitor, restore and maintain energy balance. Homeostatic signals can also be overridden, causing us to engage in non-homeostatic eating that is driven by the sensory and/or hedonic properties of palatable foods with high fat and/or sugar contents. This ‘hedonic’ eating activates brain regions involved in reward, resulting in increased motivation to consume food, and increased energy consumption [5–7] despite a state of satiety [8].

The ‘incentive salience theory’ of reward has provided a valuable framework for more than a decade for investigating the role of food hedonics in the context of appetitive behavior in humans and animals [9, 10]. This theory postulates that reward is not a unitary process, but comprises an affective pleasure component referred to as ‘liking’ and a non-affective motivational component referred to as ‘wanting’. ‘Liking’ is the hedonic component that reflects the immediate experience of eating a pleasurable food [11]. ‘Wanting’ is the incentive motivation (reward-seeking) that can lead to increased appetite, food cravings and also to overconsumption of food [9, 12, 13]. Looking at these psychological traits in eating behaviors [14–16], overeating in some individuals might reflect the abnormal functioning of either the ‘wanting’ or ‘liking’ mechanisms, and there are indications from subjects with eating disorders that ‘liking’ and ‘wanting’ of obesogenic foods can also be dissociated [17].

Dopaminergic neurotransmission in the nucleus accumbens (NAc) plays a key role in behaviors related to natural rewards and drugs of abuse [18–22]. In the NAc, two important cell classes are medium spiny neurons expressing D1 dopamine receptors (D1 cells) and those expressing D2 dopamine receptors (D2 cells) [23]. The precise contribution of D1 and D2 cells to reward-associated behaviors is still unresolved, and establishing a causal relationship between activation of each neuronal subtype and its effect on motivated behaviors has proven challenging. D1 cell activation is canonically considered to be related to positive rewarding events leading to persistent reinforcement, while activation of D2 cells is thought to facilitate aversion [24–26]. However, the current literature is controversial regarding the functionality of these cells and their role in motivation-related behaviors [27–30].

Therefore, increased understanding of which cells in the dopaminergic system underly brain substrates of ‘liking’ and ‘wanting’ may open an avenue to comprehending the impact of food rewards on eating behaviors linked to obesity and other eating disorders. It may also provide insight into brain circuits and mechanisms underlying both hedonic and motivational components of reward, including mechanisms common to both natural and artificial rewards.

This paper aims to further understand the differential contribution of NAc D1 and D2 cells to different components of palatable food consumption, namely ‘liking’ (hedonic/perceiving food as pleasurable) and ‘wanting’ (incentive motivational value/desire/craving). We assess ‘liking’ and ‘wanting’ from several different angles using multiple behavioral paradigms, while employing neuronal recording or optogenetic tools and analytical techniques, enabling us to better understand the forces underlying eating behavior that may lead to overconsumption.

## Results

### NAc D1 cell activity is transiently increased, while D2 cell activity is transiently increased then suppressed, during periods of free consumption of a palatable food reward

To probe the neural mechanisms underpinning reward consumption, we used *in vivo* fiber photometry to record fluorescence of cre-dependent GCaMP6s calcium indicator selectively expressed in D1 cells (in D1 receptor-cre mice) or D2 cells (in adenosine A2a receptor gene (Adora2a)-cre mice) [31] (**Figure 1a**).

**Figure 1.**
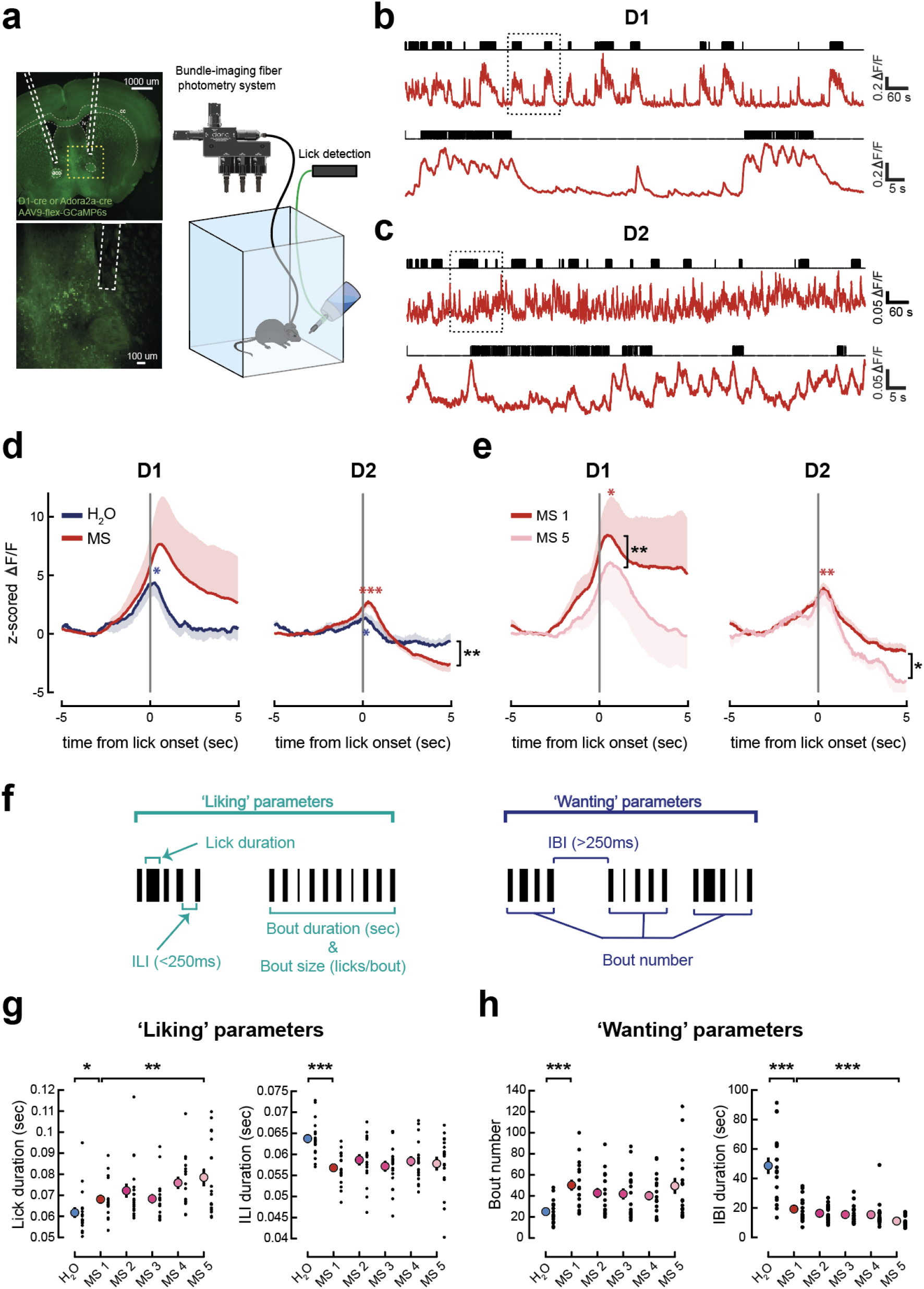
NAc D1 cells exhibit transiently elevated activity, and D2 cells exhibit suppressed activity, during periods of free consumption of palatable food. (a) Schematic of experimental set-up for photometry recordings and lick detection during an *ad libitum* consumption paradigm. Left: Illustration of fiber placement. Dashed yellow box indicates location of zoom below. (b) Top: Example photometry recording from a 20-min session with access to milkshake (MS) from a D1-GCaMP6s mouse. Licks indicated by black raster plot above. Bottom: Zoom of photometry signal and licks showing only 2 min indicated by black dotted square above. (c) As in (b) but for a D2-GCaMP6s mouse. (d) Photometry signals around water (H_2_O, blue) or MS (red) lick onset, averaged over all licking bouts in D1-GCaMP6s mice (left, n=6) and D2-GCaMP6s mice (right, n=5). MS signals are an average of 5 sessions. Mean across bouts and animals is shown, shaded area represents the SEM. D1-GCaMP6s mice, one-sample t-tests: max post lick onset water (t(5)=3.972, p=0.011), delta max-min (t(5)=8.457, p<0.001). D2-GCaMP6s mice, one-sample t-tests: max at lick onset water (t(4)=3.437, p=0.026), max at lick onset MS (t(4)=7.848, p=0.001), min post lick onset MS (t(4)=−3.526, p=0.024). Paired-samples t-test: min post lick onset MS vs. water (t(4)=5.407, p=0.006). (e) As in (d) but comparing the first and last MS sessions. D1-GCaMP6s mice, one-sample t-tests: max post lick onset MS 1 (t(5)=2.687, p=0.043), MS 5 (t(5)=1.856, N.S.). Paired-samples t-test: max MS 1 vs. MS 5 (t(5)=6.257, p=0.002). D2-GCaMP6s mice, one-sample t-tests: max post lick onset MS 1 (t(4)=7.628, p=0.002), MS 5 (t(3)=2.858, N.S.), delta max-min MS 5 (t(5)=4.274, p=0.008). Paired-samples t-test: min post lick onset MS 1 vs. MS 5 (t(3)=3.088, p=0.05). (f)Description of lick microstructure parameters associated with ‘liking’ and ‘wanting’. (g) Lick duration (left) and inter-lick interval (ILI, right) during a session with access to water (blue dot) and 5 sessions with access to MS (red and pink dots). D1- and D2-GCaMP6s cohorts pooled. Mean ± SEM values are shown in color, with individual values from each mouse overlayed in black. One-way RM-ANOVA: lick duration water vs. MS 1 (F(1,17)=4.799, p=0.043), MS 1 to MS 5 (F(4,68)=3.607, p=0.010), MS 1 vs. MS 5 (F(1,17)=9.961, p=0.006); ILI: water vs. MS 1 (F(1,17)=38.238, p<0.001), MS 1 to MS 5 (F(4,68)=0.924, N.S.), n=18. (h) As in (g) for lick bout number (left) and inter-bout interval (IBI, right). One-way RM-ANOVA: bout number water vs. MS 1 (F(1,17)=45.661, p<0.001), MS 1 to MS 5 (F(4,68)=2.268, N.S.); IBI water vs. MS 1 (F(1,17)=44.170, p<0.001), MS 1 to MS 5 (F(4,68)=5.318, p=0.001), MS 1 vs. MS 5 (F(1,17)=35.626, p<0.001), n=18. IBI: Inter-bout interval, ILI: inter-lick interval, MS: Milkshake. p-values reported on the figures as follows: *p≤0.05, **p<0.01, ***p<0.001

To determine how D1 and D2 cells represented solutions with different palatability, mice were first given free access to water from a bottle (**Figure S1a,b**). Once they reliably consumed this, they were allowed to freely consume a highly palatable food (milkshake, **Figure 1b,c**) over the course of five daily sessions, in the absence of nutrient deficit.

We found that before licking, both D1 and D2 cells transiently increased their activity, which peaked just after licking onset. In both D1 and D2 cells, the peak was more pronounced in response to milkshake than water, possibly reflecting hedonic processes (**Figure 1b-d**). However, after consumption onset, the activity patterns of the two populations differed; D1 cells returned to baseline, while D2 cell activity was suppressed below the baseline as the licking bout continued (**Figure 1d**). Both cell types continued to respond to milkshake over subsequent sessions, but the D1 peak reduced over sessions, and the D2 suppression was exaggerated (**Figure 1e**). These results demonstrate that D1 and D2 cells do not act as a single homogeneous population during consumption.

Milkshake consumption within a session (total number of licks) was significantly higher than water, and this was consistent across sessions (**Figure S1c,d**). This indicates more positive hedonic reactions to milkshake than water (**Figure S1d**). However, the amount consumed provides an incomplete readout of ingestive behavior [32]. Therefore, to further probe the driving forces behind these changes in overall consumption, we investigated different aspects of free consumption of water vs. a palatable food (milkshake). In particular, we examined if it would be possible to differentiate between ‘wanting’ and ‘liking’ parameters [14]. We therefore analyzed licking microstructure [33], since it provides several measures to evaluate pleasure and motivation during liquid-food consumption in rodents [32, 34–37]. Specifically, increases in the duration of single licks, licking bout duration and bout size (lick intensity determined by number of licks in a bout), and decreases in inter-lick interval (ILI), have been suggested to represent increased hedonic impact and palatability (‘liking’), while increases in bout number and decreases in inter-bout interval represent increased incentive salience (‘wanting’) [32, 36, 38–41](**Figure 1f**).

We found that changes in ‘liking’ parameters paralleled greater ‘liking’ for milkshake over water; lick duration, bout duration and bout size increased significantly, while inter-lick interval (ILI) decreased significantly, between the water and first milkshake sessions (**Figure 1g and S1e**). Therefore, as expected, animals displayed more positive hedonic reactions (‘liking’) to the milkshake than to water. Lick duration also kept increasing over milkshake sessions, suggesting an increased hedonic component of milkshake consumption in later sessions (**Figure 1g, left**).

Bout number increased, and inter-bout interval (IBI) decreased significantly between water and the first milkshake session, suggesting that mice were more motivated (‘wanted’) to consume milkshake (**Figure 1h**). IBI also kept decreasing across milkshake sessions, suggesting ‘wanting’ for milkshake increased over sessions (**Figure 1h, right**).

We then performed multiple regression analysis to investigate whether *in vivo* calcium activity of NAc D1 and D2 cells during consumption of milkshake across all 5 sessions could predict the level of ‘liking’ or ‘wanting’ of the palatable food consumed. This analysis can identify key features of neural activation that predict specific aspects of palatable food consumption. In D1 cells, the minimum and maximum signal values were significant predictors of bout duration (‘liking’), explaining around 30% of the variability in bout duration (**Figure S2a, top right**), with negative and positive correlations to ‘liking’, respectively. The R^2^ change values showed that the higher the initial increase (and to a smaller extent, the lower the subsequent decrease), the longer the bout (**Figure S2a**). In D2 cells, we found that the area under the curve and the minimum signal during the 3 sec following lick onset were significant predictors of bout duration, together explaining around 40% of the variability in bout duration (**Figure S2b, top right**). More specifically, the more the D2 cell signal decreases, the longer the bouts are and the higher the ‘liking’ is (**Figure S2b**). Interestingly, in a similar analysis of inter-bout interval, IBI (‘wanting’ – **Figure S3**), the significant predictors in either D1 or D2 cell signals only explained a very small amount of IBI variability and were thus poor predictors overall.

Together, these findings show that *ad libitum* consumption of milkshake induces both ‘liking’ and ‘wanting’ to eat. These two main components underlying palatable food consumption are more positively associated with consuming milkshake than water. However, we detected a dissociation between ‘liking’ and ‘wanting’, with the distinct signals in both D1 and D2 cells being more strongly correlated with ‘liking’ than ‘wanting’. Specifically, the initial activation in D1 cells at licking onset and the sustained inhibition of D2 cells activity during eating were good predictors of ‘liking’ parameters.

### NAc D2 cells increase consumption by increasing ‘wanting’ despite a decrease in ‘liking’

We next employed optogenetics to investigate how the roles of NAc D1 and D2 cells differ during *ad libitum* palatable food consumption. Based on previous findings [24, 42], we initially hypothesized that activation of NAc D1 cells would lead to increased consumption, whereas activation of NAc D2 cells – counteracting the natural suppression in activity described earlier – would decrease ‘liking’. However, it would not necessarily affect overall intake, depending on its effect on the ‘wanting’ aspect of consumption. To test our predictions, we injected the optogenetic cre-dependent excitatory actuator ChrimsonR or a control GFP virus, into the NAc of D1- or D2(A2a)-cre mice, and implanted optical fibers bilaterally in the NAc (**Figure 2a**). Consistent with our photometry data and our initial prediction, optogenetic activation of D2 cells contingent on eating decreased ‘liking’ (increased ILI, **Figure 2c**). Unexpectedly, it also increased ‘wanting’ (increased bout number, **Figure 2d**) leading to increased overall consumption (increased number of licks, **Figure 2b**). This suggests that the ‘wanting’ aspect triggered by optical activation overrode ‘liking’ in driving overall consummatory behavior. Conversely, contrary to our hypothesis, D1 cell activation did not impact overall consumption via effects on either ‘liking’ or ‘wanting’ parameters (**Figure 2b-d**). This would mean that sustained activity of D1 cells beyond the naturally occurring transient increase described in **Figure 1d, e** does not promote further ‘liking’, nor does it induce ‘wanting’.

**Figure 2.**
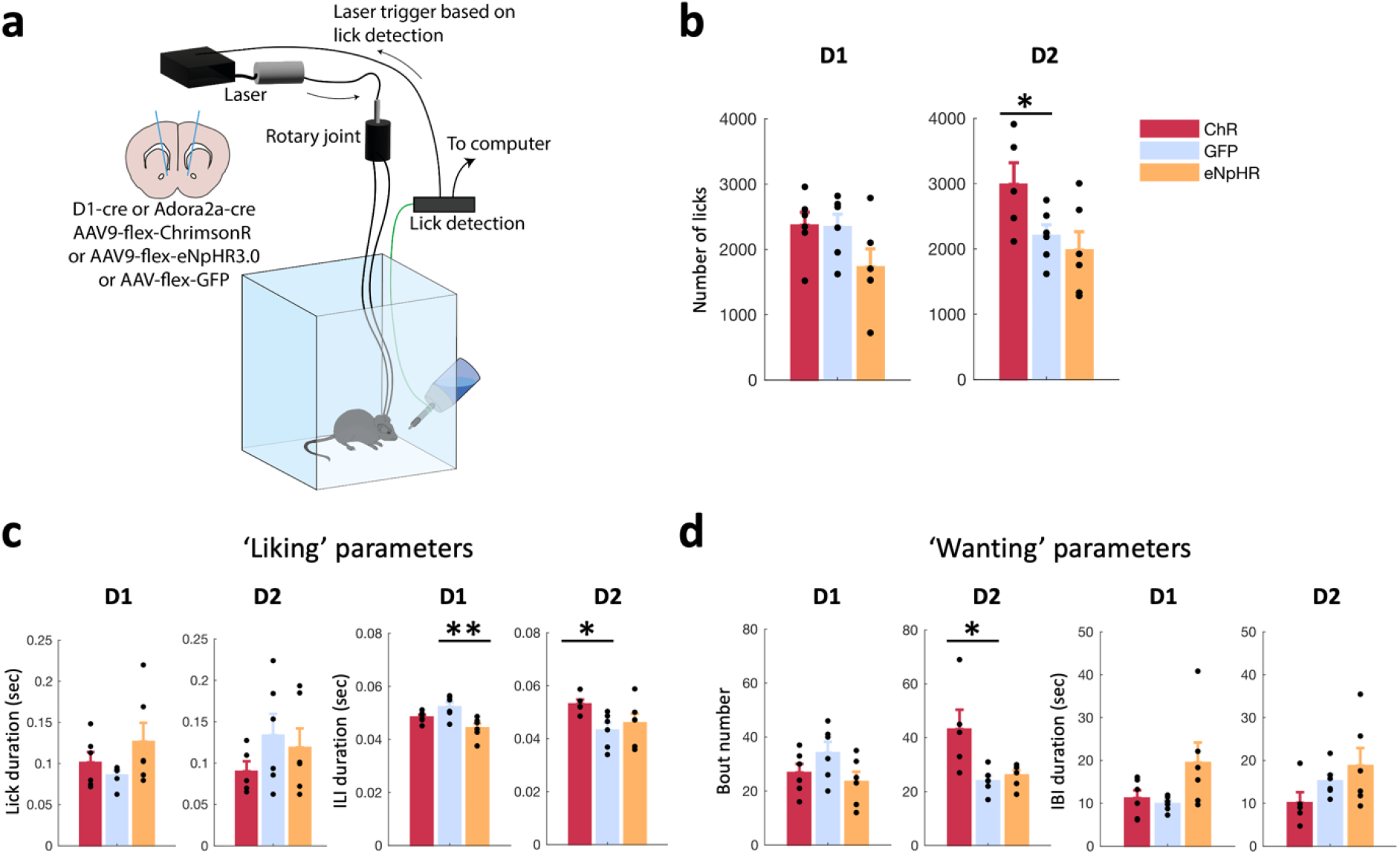
NAc D2 cell natural activity increases consumption by increasing ‘wanting’ despite a decrease in ‘liking’. (a) Schematic of experimental set-up for closed-loop optogenetic control based on recorded licks in an *ad libitum* consumption paradigm. Left: fiber placement and viruses used. (b) Total number of licks recorded in a 20-min *ad libitum* session with access to milkshake in D1-cre and D2(A2a)-cre mice, expressing either ChrimsonR (ChR, red), eNpHR3.0 (eNpHR, orange) or the GFP (blue) reporter. D1: One-way ANOVA (opsin: F(2,15)=2.564, N.S.). D2: One-way ANOVA (opsin: F(2,14)=3.837, p=0.047), independent-samples t-test: ChR vs GFP (t(9)=2.231, p=0.05). (c) ‘Liking’ parameters consisting of lick duration (left) and inter-lick interval (ILI, right) across one session with access to milkshake in D1-cre and D2(A2a)-cre mice. Mean ± SEM values are shown in color, with individual values from each mouse overlayed in black. D1: ILI, one-way ANOVA (opsin: F(2,15)=7.516, p=0.005), post-hoc independent t-test with Bonferroni correction eNpHR vs. GFP (p=0.004). D2: ILI, one-way ANOVA (opsin: F(2,14)=3.049, p=0.080), independent-samples t-test ChR vs GFP (t(9)=2.963, p=0.016). Lick duration: one-way ANOVA (opsin: D1: F(2,15)=1.813, N.S., D2: F(2,14)=0.950, N.S.). (d) As in (c) for ‘wanting’ parameters, consisting of number of licking bouts (left) and inter-bout interval (IBI, right). D1: one-way ANOVAs (opsin), Bout number (F(2,15)=2.235, N.S.), IBI: (F(2,15)=3.180, N.S.). D2: Bout number, one-way ANOVA (opsin: F(2,14)=6.642, p=0.009), post-hoc independent-samples t-test ChR vs GFP (p=0.014). IBI, one-way ANOVA (opsin: F(2,14)=2.076, N.S.). p-values reported on the figures as follows: *p≤0.05, **p<0.01.

Next, we sought to test whether silencing D1 and D2 cells using the inhibitory opsin eNpHR3.0 could also influence consumption of a palatable food. We hypothesized that D1 cell silencing would induce a decrease in consumption, while D2 cell silencing would prolong consumption. However, silencing D1 cells did not lead to a significant decrease in overall consumption (**Figure 2b**), but induced a significant increase in ‘liking’ (decreased ILI, **Figure 2c**) with no significant effect on ‘wanting’ (**Figure 2d**). The influence of inhibitory post-ingestive mechanisms such as satiation, which increase as ingestion proceeds, could have been a confounding factor to D1 cell inhibition, because the lick and bout numbers may have already been at their minimum [43, 44].

Inhibition of D2 cells did not lead to any significant difference in lick or bout number compared to controls (**Figure 2b,d)**, suggesting that a further decrease of the natural suppression in activity observed during consumption (**Figure 1d,e**) may not be possible and therefore may not yield any stronger consummatory effect.

Taken together, these results show that during *ad libitum* consumption, only D2 cell activation leads to an overall increase in eating, and this is mediated by an increase in ‘wanting’, despite a concomitant decrease in ‘liking’. However, a limitation of the *ad libitum* paradigm is that it only examines baseline motivation, and is not designed to evaluate motivation to work for a reward. Therefore, we next probed the role of D1 and D2 cells in a different motivational task, designed to interrogate ‘wanting’.

### NAc D1 and D2 cell activity increases with anticipation during a motivational task to retrieve palatable food, which is followed by a D2 cell activity decrease upon consumption onset

To probe ‘wanting’ further, we paired a food reward with a cue, which allowed us to investigate the attribution of ‘incentive salience’ to the cue. We exposed mice to a Pavlovian ‘motivation to retrieve a reward’ task in which they received food rewards announced by a sound cue, which they had to retrieve from a food magazine. This did not require the mice to develop learning strategies. This allowed us to monitor three parameters: the number of entries in the food magazine, the time to retrieve and consume the food after cue onset (latency), and the length of food-magazine entries. (**Figure 3a,b**).

**Figure 3.**
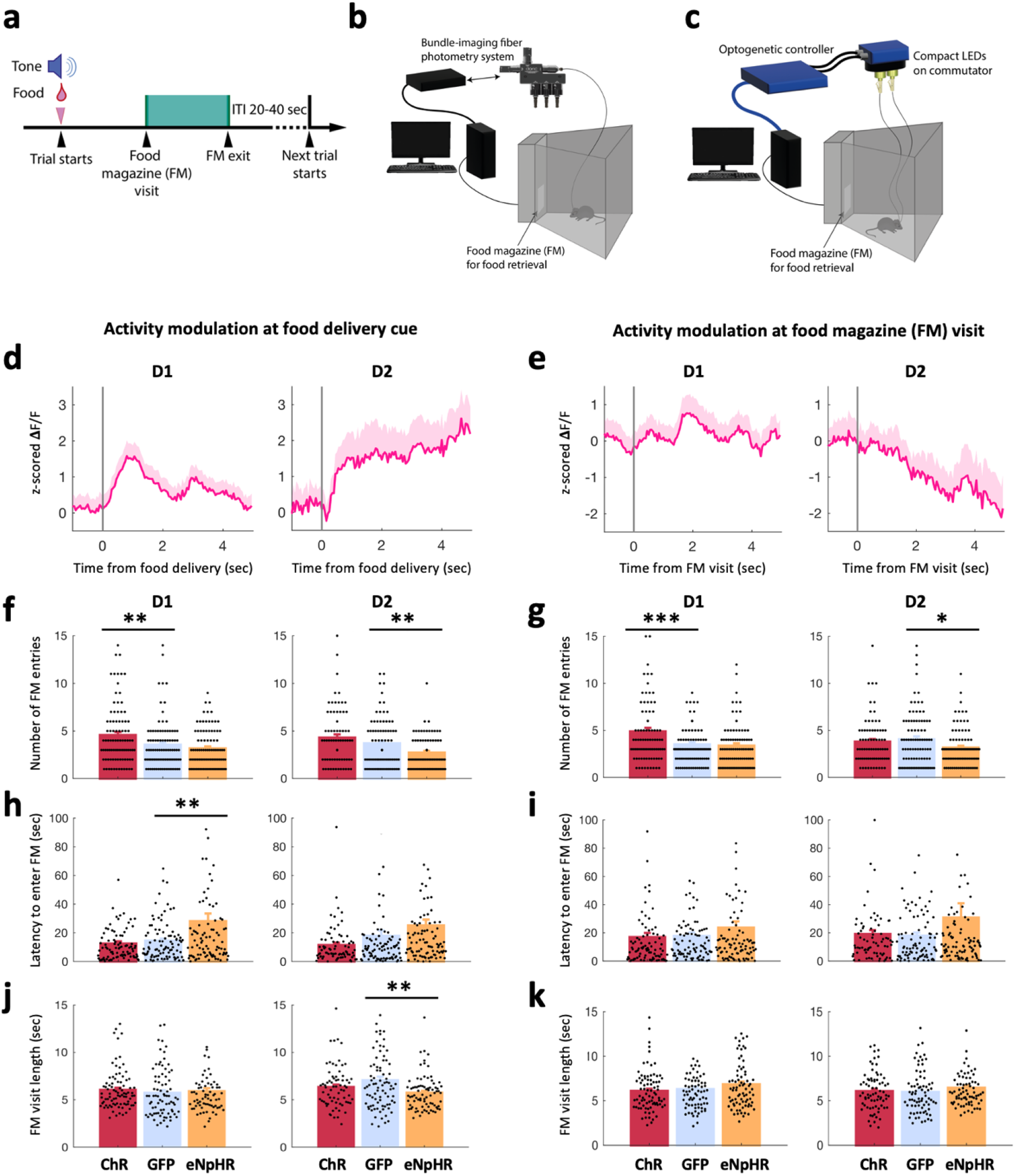
NAc D1 and D2 cell activity increases with anticipation during a motivational task and inhibition of D2 or D1 cells reduces motivation to retrieve a food reward. (a) Schematic of a food reward delivery trial structure. (b) Schematic of the chamber used for photometry recordings (applying to graphs (d) and (e)). (c) Schematic of the chamber coupled with optogenetic control triggered by the recorded behavior (applying to graphs (f)-(k), see Methods). (d) Photometry signals aligned to food delivery time, associated with a sound cue, in D1-GCaMP6s (left) and D2-GCaMP6s (right) mice. An average across mice and trials run over 2 sessions on two different days is shown. Mean across trials and animals are shown, shaded areas represent the SEM. (e) As in (d) but showing photometry signals aligned to food magazine (FM) visits. (f) Number of food magazine entries in D1-cre and D2(A2a)-cre mice expressing ChrimsonR (ChR, red), eNphR3.0 (eNpHR, orange) or GFP (blue) when a 10-sec optogenetic stimulation occurred from onset of food delivery cue. Each trial is counted as an individual data point. D1, one-way ANOVA (opsin: F(2,265)=8.630, p<0.001), post-hoc with Bonferroni correction: ChR vs. GFP (p=0.003). D2, one-way ANOVA (opsin: F(2,250)=9.828, p<0.001), post-hoc with Bonferroni correction eNpHR vs. GFP (p=0.006). (g) As in (f) but when a 10sec optogenetic stimulation occurred at the first food magazine entry of each trial. D1, one-way ANOVA (opsin: F(2,264)=9.597, p<0.001), post-hoc with Bonferroni correction ChR vs. GFP (p=0.001). D2, one-way ANOVA (opsin: F(2,252)=3.403, p<0.035), post-hoc with Bonferroni correction eNpHR vs. GFP (p=0.032). (h) As in (f) but for the latency to enter the FM following food delivery cue. Optogenetic stimulation was triggered at food delivery cue. D1, one-way ANOVA (opsin: F(2,264)=9.820, p<0.001), post-hoc with Bonferroni correction eNpHR vs. GFP (p=0.003). D2, one-way ANOVA (opsin (F(2,246)=5.158, p=0.006), post-hoc tests N.S. (i) As in (h) but when the 10-sec optogenetic stimulation occurred at the first food magazine visit of each trial. D1: one-way ANOVA (opsin: N.S.). D2: one-way ANOVA (opsin: N.S.). (j) As in (f) but showing the average length of a food magazine visit. Optogenetic stimulation occurred at food delivery cue. D1, one-way ANOVA (opsin: N.S.). D2, one-way ANOVA (opsin: F(2,209)=5.408, p=0.005), post-hoc with Bonferroni correction eNpHR vs. GFP (p=0.004). (k) As in (j) but when optogenetic stimulation occurs during 10 sec following the first FM visit of each trial. D1: one-way ANOVA (opsin: N.S.). D2: one-way ANOVA (opsin: N.S.). FM: Food magazine. p-values reported on the figures as follows: *p≤0.05, **p<0.01, ***p<0.001.

Photometry recordings showed sustained increased activity in D2 cells during the approach phase (after food delivery cue; **Figure 3d, right**). D1 cells, however, only transiently increased their activity following food delivery cue (**Figure 3d, left**). An important distinction in the behavior of D1 cell activity must be made: in contrast to the *ad libitum* paradigm where D1 cell signal increased prior to licking onset, here, in a motivational task, the increase in signal was specifically aligned to the reward delivery cue. During food magazine visits, the neural dynamics observed were similar to the first *ad libitum* study, with a sustained decrease in the activity of D2 cells and only a small initial increase in D1 cells (**Figure 3e**).

As in the first experiment, we were interested to see if the natural D1 and D2 cell activity held predictive information about components of feeding behavior. Correlation analysis of ‘wanting’ behavioral parameters and D2 cell activity after a food delivery cue revealed that latency to enter food magazine can be well predicted by the average D2 cell activity before the food magazine visit, and by the standard deviation (Std) of the pre-food magazine visit D2 cell signal (**Figure S4a;** the higher the signal before visit, the shorter the latency). This correlation analysis also showed that the number of food magazine visits can be well predicted by the median D2 cell signal before the food magazine visit (**Figure S4b;** the higher the signal pre-food magazine visit, the more visits occurred). The average length of a food magazine visit was also found to be well predicted by the minimum of D2 cell activity *during* consumption, and by the standard deviation of the signal before the visit (**Figure S4c**; the lower the minimum, the longer the food magazine visit). D1 cell analysis yielded no significant correlations.

Therefore, latency and number of visits correlated with D2 cell signals *before* visits occurred, while the length of a visit was determined by the signals *during* the visit. These data also suggest that suppression of D2 cell activity *during* feeding favors consumption, which was investigated further in the next experiment.

### Inhibition of D2 or D1 cells reduces motivation to retrieve a food reward, while activation of D1 cells, but not D2 cells, enhances motivation

Because both D1 and D2 cells were modulated during the motivation task (**Figure 3d,e**), we aimed to causally test the effect of cell-type-specific activation and inhibition on motivated behavior (‘wanting’). We optogenetically manipulated NAc D1 and D2 cells at either food reward delivery cue or at food magazine visit (first visit following a delivery cue) (**Figure 3a,c**). D2 cell silencing at either food delivery cue, or at food magazine visit significantly decreased number of food magazine entries (**Figure 3f,g**), and inhibiting D2 cells after food delivery cue significantly decreased the average length of the subsequent food magazine visit (**Figure 3j**). These results are in line with our photometry findings (**Figure 3d, right**) suggesting a role for D2 cell activity in the anticipation of a food reward (‘wanting’). Optogenetic activation of D1 cells at either food delivery cue or at food magazine visit significantly increased the number of food magazine entries (**Figure 3f,g**). Moreover, silencing D1 cells during the food delivery cue increased the average latency to food magazine entry (**Figure 3h**). Combined with photometry findings (**Figure 3d,e, left**), these findings about D1 cells suggest that their activity causes a higher incentive motivation to approach the palatable food, making the mice more focused on obtaining the reward and more reactive to the cue.

Together, our data indicate that both D1 and D2 cell activity during the food approach code and control incentive motivation or ‘wanting’.

### Modulation of NAc D1 but not D2 cells influences hedonic preference learning (hedonic shifting)

Our two previous experimental paradigms showed that consumption of a palatable food elicits, as expected, ‘liking’ responses. We next investigated whether pairing a flavored palatable food, constituting a conditioned stimulus (CS), with optogenetic stimulation (unconditioned stimulus, US) of either D1 or D2 cells could shift natural preference for one flavor to another. We hypothesized that optogenetic stimulation of either population would establish a learned preference for the least innately preferred flavor by increasing the hedonic value (‘liking’) of the CS (flavored food).

To assess innate preference, mice were first presented with two bottles, each containing a different flavor, but with identical nutritional composition (‘pre-test’). The least preferred flavor was then coupled to optogenetic stimulation of either D1 or D2 cells during a daily conditioning session over several days, while the preferred flavor was presented unpaired with optogenetic stimulation in a separate daily session (**Figure 4a**). Following conditioning, we performed a ‘choice-test’ to determine if the paired flavor was now able to trigger incentive motivation (‘wanting’) and become equally, or possibly more, attractive than the previously preferred, unpaired flavor. During conditioning, increased consumption from the paired flavor over the unpaired one would indicate a hedonic change (‘liking’). During the choice test, with respect to the ‘liking’/‘wanting’ hypothesis, disruption of a reward-oriented behavior by optogenetic manipulation of the NAc could, in principle, be due to a disruption of either ‘liking’ or ‘wanting’ or, presumably, a combination of both.

**Figure 4.**
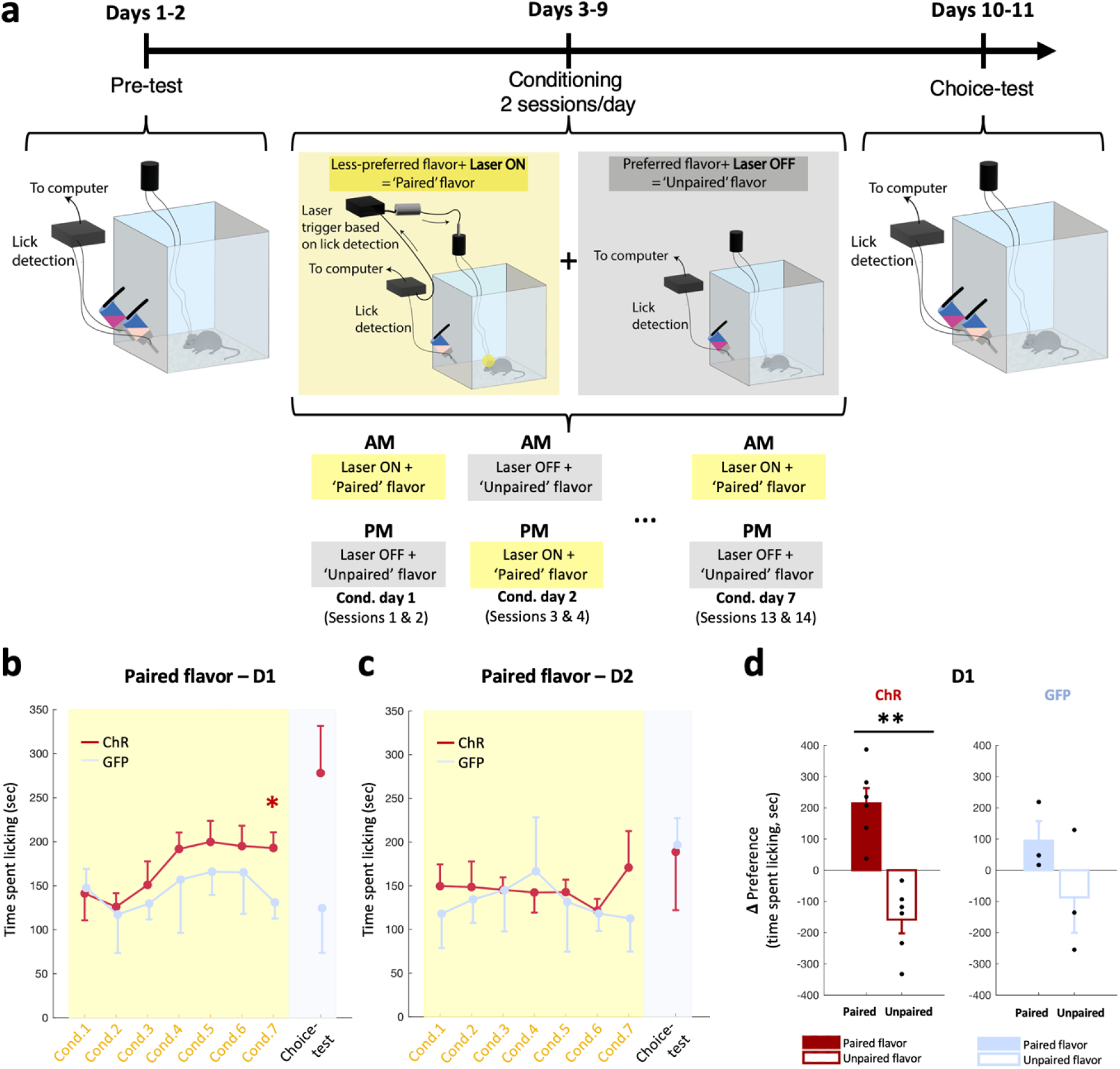
Modulation of NAc D1 and not D2 cells influence hedonic preference learning (hedonic shifting). (a) Schematic of the experimental design for lick detection and closed-loop optogenetic activation on the paired-flavor. Left, during the pre-test, mice had free access to both flavored bottles. Middle, mice were given “Laser ON” sessions (yellow box), where only the initially less preferred flavor was presented, and licking was paired with opto-stimulation. On the same day, the initially more preferred flavor was given and the laser was not triggered. Sequence of laser ON and OFF conditioning sessions counterbalanced from day to day (bottom). Right, in the ‘choice test’, the two flavors were once again presented together, in the absence of optostimulation. (b) Time spent licking on the paired-flavor spout during conditioning sessions (Cond., yellow box) and last choice-test session (grey box) in D1-cre mice expressing ChrimsonR (ChR, red) and GFP (blue). One-sided independent-samples t-test: ChR vs. GFP (Cond.7: t(7)=2.160, p=0.034; Choice-test: t(7)=1.804, p=0.057). (c) As in (b) but in D2(A2a)-cre mice. Mixed-design ANOVA: N.S. (d) Change in preference expressed as the difference (**Δ**) in time spent licking between the choice-test and the pre-test (choice-test – pre-test) on a given spout for the paired (filled bars) and unpaired (empty bars) in D1-ChR (left, red) and D1-GFP (right, blue) mice. ChR, paired-samples t-test paired vs. unpaired flavors: t(5)=4.754, p=0.003 – one-sided. GFP: t(2)=1.074, p=0.198 – one-sided. ChR: ChrimsonR, Cond.: Conditioning. p-values reported on the figures as follows: *p≤0.05, **p<0.01.

During optogenetic activation of D2 cells, consumption of the paired and unpaired flavors across the conditioning phase did not differ (**Figure 4c**). Note that during this conditioning phase, each flavor was presented in separate sessions, hence the absence of difference between the amounts consumed, unlike the difference observed in the ‘pre-test’ (**Figure S5a, right**). However, optogenetic stimulation of D1 cells led to a gradual increase in the consumption of the paired flavor, so that the mice consumed larger amounts of food in the late paired conditioning sessions compared to GFP controls (**Figure 4b**). During the choice-test, an increase in consumption of the paired flavor compared to the pre-test levels was noted while a decrease was observed in the unpaired flavor, leading to significantly different ‘choice-test – pre-test’ deltas between the two flavors (**Figure 4d, left**). There was no significant change in preference in control GFP mice (**Figure 4d, right**), suggesting repeated exposure to the flavors did not alter preference.

In addition, we used lick microstructure analysis to assess potential changes in ‘liking’ and ‘wanting’ produced by flavor preference conditioning. During the conditioning sessions, stimulation of D1, but not D2 cells increased ‘liking’ (increased lick duration and decreased inter-lick duration) over time (**Figure 5a,b, left**). This indicates an increased palatability of the flavor paired with D1 cell optostimulation, which is of relevance because conditioned increases in palatability have been shown to be important for driving overeating [45]. Moreover, inter-bout interval (IBI) decreased over conditioning sessions (**Figure 5d, left**), indicating an increase in ‘wanting’, while motivation to consume the unpaired flavor decreased over sessions, as indicated by decreasing bout numbers and initially increasing IBI (**Figure 5c,d, middle**). During the choice-test, when both flavors were once again presented simultaneously, with no optogenetic stimulation on either of the flavors at this stage), licking microstructure indicated an increased ‘liking’ of the paired flavor (higher lick duration, and lower ILI, **Figure 5a,b, right**). These effects were not observable in the unpaired flavor. ‘Wanting’ was also increased for the paired flavor (increased bout number, decreased IBI, **Figure 5c,d, right**) and decreased for the unpaired flavor (**Figure 5d, right**).

**Figure 5.**
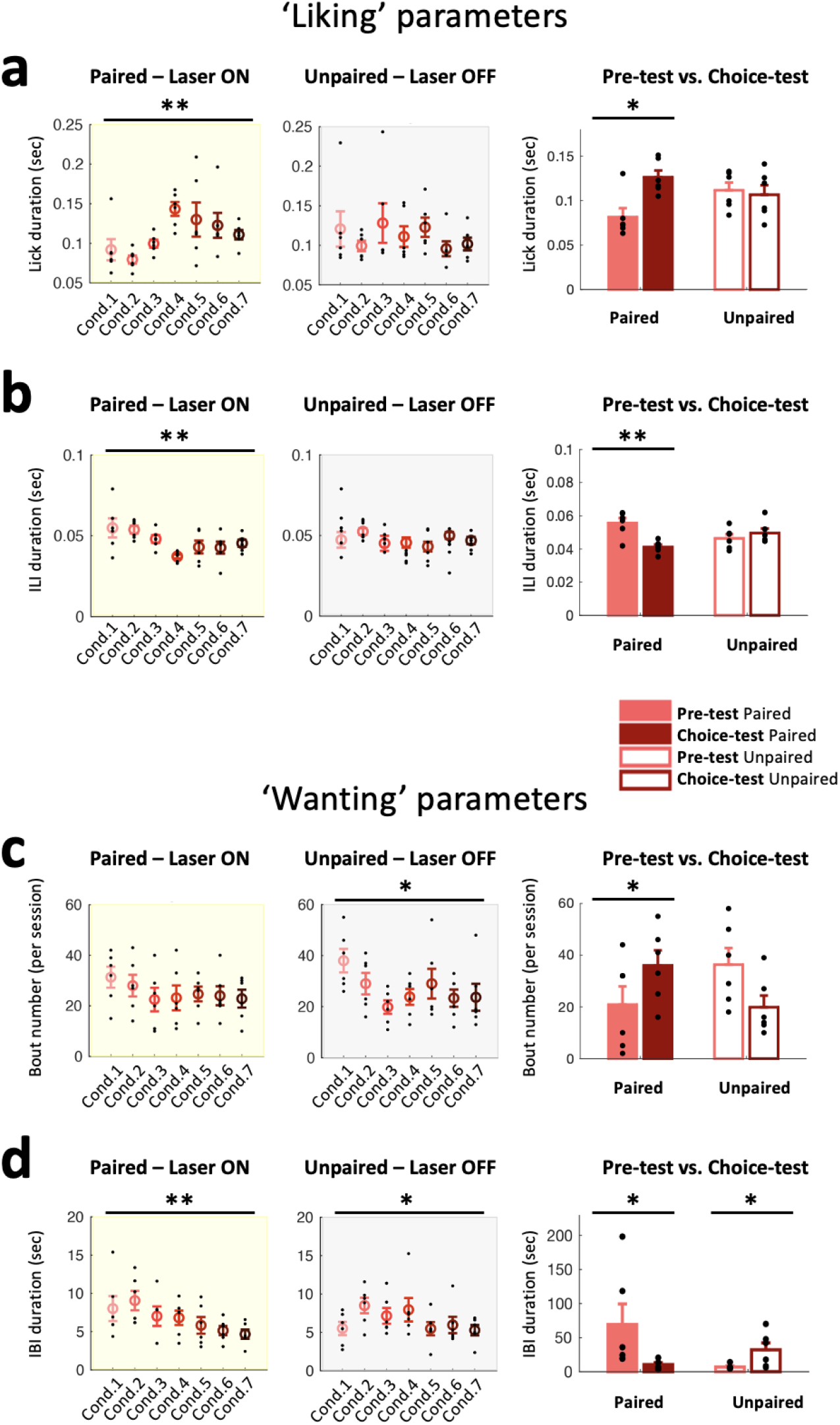
Activation of NAc D1 cells induces a preference shift via an increase in ‘liking’ and ‘wanting’ for the optoactivation-paired flavor. (a) Single-lick duration across conditioning sessions (Cond.) for laser ON (left, yellow box) and laser OFF (middle, grey box) and pre-test & choice-test sessions (right) for laser-paired (filled bars) and laser-unpaired (unfilled bars) flavors in D1-ChR mice. Mean ± SEM are plotted in color, and individual values from each mouse in black. Conditioning, paired flavor: RM-ANOVA (day: F(6,30)=3.386, p=0.006 – one-sided), one-sided paired-sample t-test Cond.1 vs. Cond.3: t(5)=−2.675, p=0.022; unpaired flavor (day: F(6,30)=0.825, N.S.). Choice-test vs. pre-test, paired flavor: one-sided paired-sample t-test, t(5)=−3.188, p=0.012; unpaired flavor, t(5)=0.310, N.S. (b) As in (a) but for inter-lick interval (ILI) durations. Conditioning, paired flavor: RM-ANOVA (day: F(6,30)=3.870, p=0.003 – one-sided), Cond.1 vs. Cond.4 (one-sided paired-sample t-test, t(5)=3.212, p=0.012); unpaired flavor (day: F(6,30)=1.334, N.S.). Choice-test vs. pre-test, paired flavor: one-sided paired-sample t-test, t(5)=4.422, p= 0.003; unpaired flavor: t(5)=−0.834, N.S. (c) As in (a) but for the number of licking bouts. Conditioning, paired flavor: RM-ANOVA (day: F(6,30)=0.872, N.S.); unpaired flavor (day: F(6,30)=2.588, p=0.019 – one-sided), Cond.1 vs. Cond.7 (one-sided paired-samples t-test, t(5)=2.330, p=0.034). Choice-test vs. pre-test, paired flavor: one-sided paired-samples t-test, t(5)=−1.986, p=0.05; unpaired flavor, t(5)=1.800, p=0.066. (d) As in (a) but for the inter-bout interval (IBI) durations. Conditioning, paired flavor: RM-ANOVA (day: F(6,30)=3.176, p=0.008 – one-sided), Cond.1 vs. Cond.7 (one-sided paired-samples t-test, t(5)=2.074, p=0.046); unpaired flavor (day: F(6,30)=2.280, p=0.031 – one-sided), Cond.1 vs. Cond.7 (one-sided paired-samples t-test, t(5)=0.276, p=0.397), Cond.1 vs. Cond.2 (t(5)=−7.664, p<0.001). Choice-test vs. pre-test, paired flavor: one-sided paired-samples t-test, t(5)=1.961, p=0.05; unpaired flavor: t(5)=−2.471, p=0.028. ChR: ChrimsonR, Cond.: Conditioning, IBI: inter-bout interval, ILI: inter-lick interval. p-values reported on the figures as follows: *p≤0.05, **p<0.01.

These results show an overall preference for the flavor coupled with D1 cell stimulation, thus demonstrating an important role for D1 cells in promoting the consumption of preferred palatable foods.

## Discussion

Our study reveals distinct patterns of NAc D1 and D2 cell activity during different behavioral paradigms intended to assess different psychological processes of hedonic behaviors: hedonia (‘liking’ a food) and motivation (‘wanting’ the food). We found that D1 and D2 cells do not act as a single homogeneous population in regulating these processes (**Figure 6**). In a free palatable food consumption paradigm, a dual role of D2 cells is linked to both ‘liking’ and ‘wanting’: increased D2 cell activity signals ‘wanting’ and decreased activity enhances ‘liking’, with the magnitude of natural inhibition following palatable food consumption being a good predictor for ‘liking’. In contrast, D1 cells promote ‘liking’ during the free consumption. In the operant task, we showed that D1 cell activity after cue and right after eating onset drives the motivation to retrieve palatable food, while reduced activity of D2 cells after food magazine visits drives the motivation to remain in the magazine. Lastly, we show that D1 cell stimulation induced hedonic shifting in a Pavlovian task, leading to overconsumption of palatable food. This effect was specific to D1 cells, as stimulation of D2 cells during opto-paired food conditioning had no effect across conditioning days or during the choice test.

**Figure 6.**
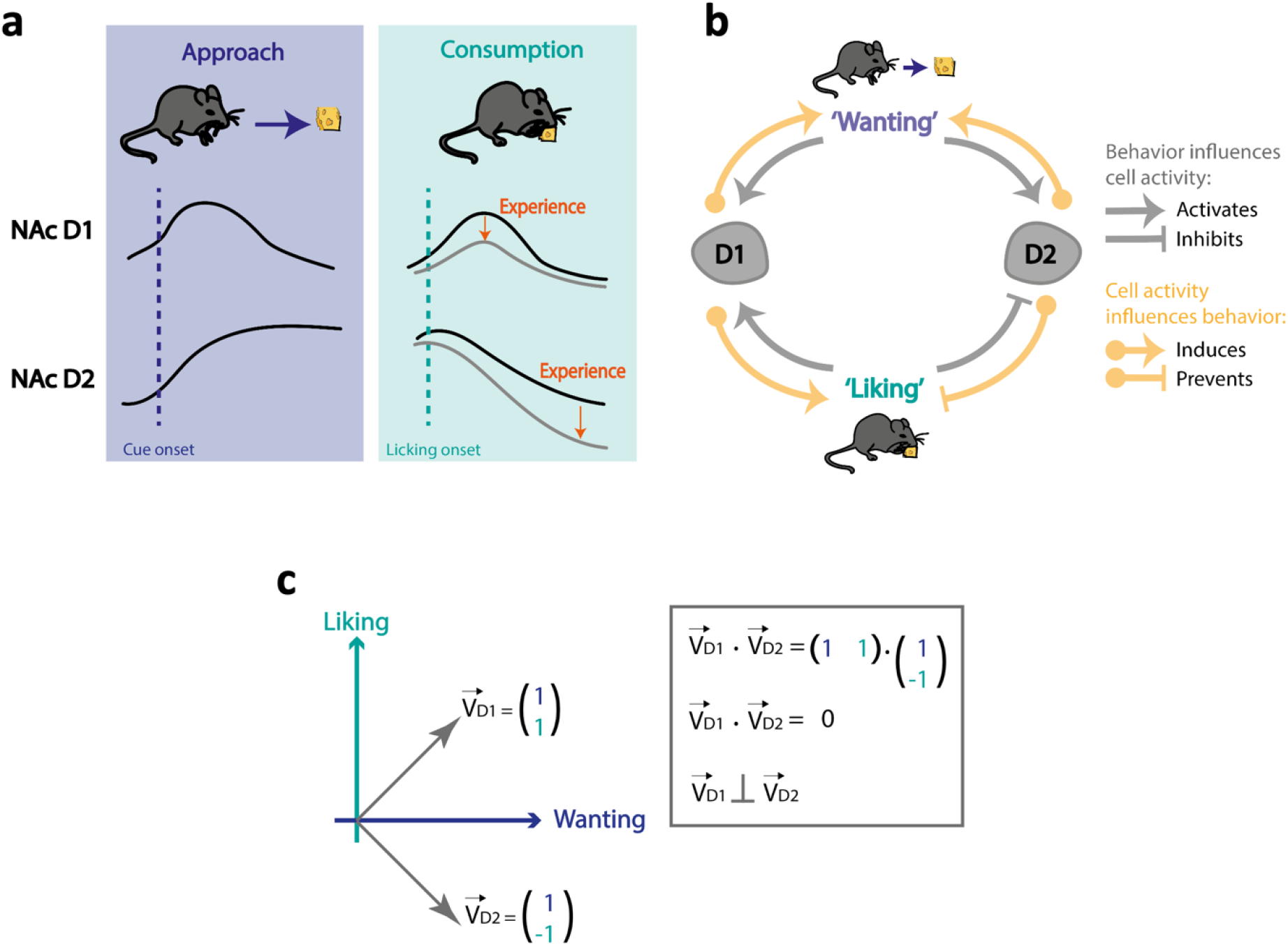
NAc D1 cells positively induce ‘wanting’ and ‘liking’ while D2 cells positively encode ‘wanting’ but negatively induce ‘liking’. Summary of the findings from the different experimental paradigms to test the distinctive patterns of NAc D1 and D2 cell activity relative to distinct psychological processes of hedonic behaviors: ‘liking’ (hedonia) and ‘wanting’ (motivation). (a) Summary of D1 and D2 cell activity in the NAc around food approach (left, blue box) and following food consumption onset (right, turquoise box), and the effect of repeated exposure to the reward. NAc D1 cells show transient activation following a reward-predicting cue (left, blue dotted line) and following consumption onset (right, turquoise dotted line). The amplitude of increase following feeding onset diminishes with daily repeated exposure to the food reward (grey line and orange arrow, see **Figures 3d,e**). NAc D2 cells show a sustained increase in activity following a food-predicting cue and their activity drops following feeding onset; this drop becomes lower during the consumption bout and with repeated daily exposure to the food reward (orange arrow). (b) Hypothetical model of the mutual relationship between approaching (‘wanting’) and consuming (‘liking’) reward behaviors, and NAc D1 and D2 cell activity. NAc D1 cell activation induces both ‘liking’ and ‘wanting’ processes (yellow arrows), and D1 cells are naturally activated during behaviors associated with ‘wanting’ and/or ‘liking’ (grey arrows). NAc D2 cell activation, on the other hand, leads to increased ‘wanting’ and decreased ‘liking’ (yellow arrows). Behaviors associated with ‘wanting’, e.g., following a food-predicting cue and during food approach, lead to an increase in D2 cell activity, while behaviors associated with ‘liking’, e.g., during food consumption, lead to a sustained decrease in D2 cell activity in the NAc. (c) Considering ‘liking’ and ‘wanting’ as a 2D space underlying hedonic eating, our data show that D1 and D2 cells, if expressed in a binary manner, are orthogonal in that space and can themselves form a base of this 2D space. This would support the idea that evolutionary processes led to these two cell populations that can independently and mutually influence hedonic eating. D1 cells are prone to lead to a maladaptive cycle of increased ‘liking’ and ‘wanting’, thus driving overeating, while D2 cells cannot reinforce both processes and are faced with a tradeoff between increasing ‘wanting’ by being more active or allowing ‘liking’ by remaining silent.

### The role of NAc D1 and D2 cells during periods of free consumption of palatable food

The NAc is part of a neural circuit governing consumption of palatable food [46] and neuronal subpopulations in the NAc shell encode rewarding stimuli and palatability [47, 48]. Pharmacological manipulations in rodents and in hyper-dopaminergic knockout mice showed that ‘wanting’ and ‘liking’ sucrose could be separated with an increase in ‘wanting’ but a constant decrease in ‘liking’ [49–52] and that ‘wanting’ was increased with prolonged sucrose consumption [53]. Here, using photometry and optogenetics, we showed a differential role between NAc D1 and D2 cells during free consumption of a palatable food. Activity of NAc D2 cells was linked to both ‘liking’ and ‘wanting’ but with opposing actions; high D2 cell activity signaled ‘wanting’, and low D2 cell activity enhanced ‘liking’. Importantly, the increased activity in D2 cells was similar across milkshake sessions, whereas inhibition of this signaling following the onset of milkshake consumption was greater after repeated milkshake sessions (**Figure 1e**). A multiple regression analysis indicated that this magnitude of inhibition (area under the curve) after consumption was a good predictor for ‘liking’ (bout duration, **Figure S2b**). Previous findings showed that during consumption of sucrose, a decrease in firing rate in a subset of NAc neurons immediately before and during a lick bout is required to initiate and maintain consummatory behavior [54, 55]. The present study supports the idea that sustained D2 cell depression maintains the actions resulting in palatable food consumption and hedonic effects [47, 54–57]. We showed that D1 cells on the other hand promote ‘liking’ right after consumption onset. D1 receptors have been shown to play a role in incentive learning [58], and so may be important during the initial phase of instrumental learning, when reward is novel and unpredictable [59] and in the primary response to drugs of abuse [60]. This may explain why D1 cell activity was highest during the first milkshake session (**Figure 1e**). Supporting this interpretation, in a multiple regression analysis, the maximum D1 cell signal following lick onset was a significant predictor of bout duration; the larger the initial increase, the longer the licking bout (**Figure S2a**).

### Contribution of NAc D1 and D2 cells to reward-associated cues

D1 and D2 cells displayed distinct temporal activity during food magazine visits in the motivational task; D1 cell activity increased after onset of the food delivery cue, whereas D2 cell activity was increased before, but suppressed after, food magazine entry (**Figure 3d,e**). These results are in line with previous findings from drug-addiction literature showing that NAc D1 cells increase their activity before entering a drug-paired chamber, while D2 cells decrease their activity after [61]. Thus it may be that in our task, D1 cell activity drives the motivation to enter the food magazine, while reduced activity of D2 cells after food magazine visit onset would drive the motivation to remain in the magazine. This might suggest that D1, but not D2 cell signaling drives reward- and reinforcement-related behaviors. In particular, we show a causal role of temporally specific cue-elicited D1 cell signaling, which may further demonstrate the role of this cell type in motivation to retrieve rewards. Activation of D1 cells when animals entered the food magazine or at cue-onset increased the number of food magazine visits. This may imply that the D1 cell signals guide animals to seek reward, whereas suppressing D1 cell activity at either food magazine or cue onset does not lead to any changes in the number of visits compared to the control group. However, inhibition of D1 cells at cue onset delayed approach. D1 cell activity may thus drive higher incentive motivation to reach the palatable food (reward) by increasing focus on the cue, and amplifying ‘wanting’ for a particular incentive target (the highly-caloric palatable food). This could contribute to intense urges to indulge in those foods, leading to overeating. Attenuating or eliminating D1 cell signals may decrease the associative learning of environmental stimuli (reward-associated cue) with reward. Our experiments were conducted once the animals learnt the association between cue and reward and the neuronal response that was measured was to the reward-predictive cue [62, 63]. Pharmacological studies investigating the role of NAc D1 and D2 receptors in response to drugs of abuse have also shown that both D1 and D2 receptors have a role in associative learning [64]. Dopamine responses to cues predicting drug availability can lead to drug seeking behavior and over time these responses to drug-associated cues are dampened [60, 64]. Our findings regarding D1 cells in the context of associative learning are in line with previous publications demonstrating that D1-like receptors are essential in reward related learning including instrumental learning [65, 66] as well as driving motivation to action [67]. More specifically, systemic administration as well as local infusion into the NAc of dopamine D1-like receptor antagonists have been shown to attenuate food-reinforced lever pressing and to dampen the rewarding effects of palatable food [68–72]. Furthermore, optogenetic activation of NAc D1 cells enhanced drug-induced conditioned place preference [24, 73]. Pharmacological activation of NAc D1 receptors produced similar findings, showing that D1 cell activity drives incentive motivation to consume food and drugs of abuse [30, 74, 75].

On the other hand, D2 cell silencing at the time of food delivery cue and at the time of food magazine visit significantly decreased the number of food magazine entries. Also, silencing natural D2 cell activity after the food delivery cue also significantly decreased the average length of a subsequent food magazine visit. These observations suggest a role for D2 cell activity in the anticipation of a reward (‘wanting’). These effects on motivation, although inconsistent with some previous findings which suggested that D2 cells are associated with negative rather than positive valence events [1, 24, 42], were in line with other more recent findings which have shown that D2 cell activation increased motivation to obtain a reward in a progressive ratio task [22, 76].

### NAc D1 cell stimulation drives hedonic shifting and overconsumption of palatable food

In the hedonic-shifting choice test after the final optogenetic stimulation pairing, D1 but not D2 cell activity significantly increased preference for the flavor that was previously least preferred. This indicates that D1 cell optostimulation induced a hedonic preference via an association between the flavor and the rewarding effects of stimulation (**Figure 4d**). This was not observed in control mice. In addition, during the conditioning sessions with D1 cell optostimulation, an increase in time spent consuming the opto-paired flavor was observed across sessions (**Figure 4b**). This may reflect a hedonic experience which alters associative hedonic/‘liking’ processes by selectively enhancing D1 cell activity. This facilitates the association between flavor and opto-stimulation and promotes hedonic shifting that in turn leads to increased consumption of that flavor as observed in the choice test. This effect was specific for NAc D1 cells; stimulation of NAc D2 cells during conditioning had no effect either across conditioning days, or on the choice test.

In this experiment we were able to induce a learned shift in preference for a particular flavor when D1 cell optogenetic stimulation was coupled with a less preferred flavor. This is based on a Pavlovian association formed between NAc D1 cell activation (US) and the least preferred flavor (CS), which changed the taste preference to that flavor. This suggests that hedonic ‘liking’ of a flavor stimulus can be determined not only by flavor itself, but also by the relevant ‘brain’ state at the time of formation of Pavlovian associations, for example by a change in perceived palatability. In a similar manner, in the context of drug addiction, it was shown that NAc D1 cell activity preceding entry to the drug-paired compartment in a conditioned place preference test is needed for drug-associative learning to take place [24, 61].

That our findings are not always in line with previous literature may reflect the discordance between previous studies themselves: the precise NAc subregion targeted plays a role in the results obtained [27, 77], the pattern (frequency, length) of the optogenetic modulation pattern also has an impact on the direction of the effects [29], and the experimental paradigm also plays a crucial role depending on the cues and responses it entails [78].

### Clinical implications for the role of NAc D1 and D2 cell activity in ‘liking’ versus ‘wanting’ in overconsumption and eating disorders

Our findings have potential implications for human eating disorders. ‘Hedonic eating’ does not necessarily involve pleasure, but can also be driven by a motivation to eat which can be dissociated from the drive caused by a nutrient deficit [79, 80]. It is already well known that in humans, high-calorie foods, especially sweet and fatty ones, promote overconsumption. The incentive theory of ‘liking’/‘wanting’ is significant in the study of food intake regulation, modelling associations between the function of the reward system and feeding or obesity. Studies tried to show that different neural mechanisms are responsible for ‘wanting’ a certain food than ‘liking’ it. For more than a decade, human studies of obesity have used ‘liking’ and ‘wanting’ to try to better understand how ‘wanting’ food can differ in some individuals, and override ‘liking’ to cause excessive cue-trigged ‘wanting’ (craving) that will drive overeating and lead to obesity [81–85]. Incentive sensitization theory, originally postulated for drug addiction [86], was then applied to eating disorders to try to explain how some individuals may experience ‘wanting’ to eat palatable foods [14, 15]. Neuroimaging studies suggest that in humans diagnosed with obesity or binge-eating disorders, overeating is triggered by visual cues associated with palatable food [87–89]. For example, increased brain activity in response to palatable foods was positively correlated with self-reported craving rates or ‘wanting’ to eat compared to healthy controls [90–92]. This evidence is very similar to findings in individuals suffering from drug addiction [83, 93–97].

### Conclusion

To our knowledge, our study is the first to approach aspects of NAc dependent reward consumption using such a multi-faceted approach. We incorporated observation, inhibition and excitation of NAc D1 and D2 cells, multiple behavioral paradigms investigating different aspects of ‘liking’ and ‘wanting’, including analysis of lick microstructure, and multivariate analysis. In summary, we show that NAc D1 cell activity plays a role in reinforced behaviors induced by palatable foods in an operant paradigm or a Pavlovian association test. However, in a palatable food free-access paradigm D1 cells encode only the hedonic property of the palatable food. NAc D2 cells on the other hand, have a role in the drive to retrieve food, showing elevated activity prior to eating onset in a motivational task. Once eating starts, suppressed D2 cell activity allows for prolonged eating episodes. This would support the idea that evolutionary processes led to these two cell populations that can independently and mutually influence hedonic eating. D1 cells are prone to lead to a maladaptive cycle of increased ‘liking’ and ‘wanting’, thus driving overeating, while D2 cells cannot reinforce both processes and are faced with a tradeoff between increasing ‘wanting’ by being more active or allowing ‘liking’ by remaining silent (**Figure 6**). These findings, which temporally distinguish between the different psychological processes of food reward and reinforcement, support the incentive salience hypothesis [9, 15]. They may assist the design of future studies and improve diagnostic criteria of addiction and eating disorders.

## Materials and Methods

### Animals and housing conditions

All animal procedures were carried out in accordance with the Animal Welfare Ordinance (TSchV 455.1) of the Swiss Federal Food Safety and Veterinary Office and were approved by the Zurich Cantonal Veterinary Office. Mice were kept on a reversed 12-h/12-h light/dark cycle and provided with standard chow pellets *ad libitum*. In order to avoid food deprivation for behavioral testing, mice were provided with 2% citric acid in water *ad libitum* in their home cage [98]. All experiments were performed during the dark phase. Both adult male and female mice were used for the experiments (see Table 1) and mice were at least 7 weeks old before surgeries were performed. As no differences were found in the results between sexes, males and females were pooled (see Statistical Analysis section).

**Table 1:** Mice implanted in each group and line.

|  | Adora2a-cre | D1-cre |
| --- | --- | --- |
| *Photometry – GCaMP6s* | 6 (4m + 2f) | 6 (3m + 3f) |
| *Optogenetics – ChrimsonR* | 5 (3m + 2f) | 6 (3m + 3f) |
| *Optogenetics – eNpHR3.0* | 6 (3m + 3f) | 6 (3m + 3f) |
| *Optogenetics – GFP* | 6 (3m + 3f) | 6 (3m + 3f) |

### Viruses and surgeries

For specific targeting of D1 cells and D2 cells, D1-cre (B6.FVB(Cg)-Tg(Drd1-cre)EY262Gsat/Mmucd) and Adora2a-cre (Adora2aTg(Adora2a-cre)KG139Gsat, originally obtained from the Jackson Laboratory) mouse lines were used, respectively. For photometry recordings, 6 mice from each line were injected bilaterally with a pAAV9-CAG-Flex.GCaMP6s.WPRE.SV40 virus (Addgene, titre ≥ 1×10^13^ vg/mL) in the medial NAc shell (D1-cre and D2(A2a)-cre mice; bregma +1.4mm, midline ± 1.35mm, depth 3.9 mm, at an angle of 10°) and were then implanted bilaterally with optic fibers (200 μm diameter, NA 0.37, Doric Lenses), using the same coordinates (except that the depth of the fiber tip was reduced to 3.8mm).

For optogenetic modulation, mice (see Table 1) were injected with pAAV9-EF1a-DIO-ChrimsonR-mRuby2-KV2.1-WPRE-8V40 (Addgene, titer: 1×10^13^ vg/mL) or ssAAV-9/2-hEF1a-dlox-eNpHR3.0_iRFP(rev)-dlox-WPRE-hGHp(A) (ETHZ/UZH Viral Vector Facility, titre: 7.2 × 10^12^ vg/mL) or pAAV-flex-GFP (Addgene, titer: 2.5 ×10^13^ vg/mL) for activation, inhibition and control respectively. Viruses and fiber implants were positioned at the same coordinates as the photometry cohort. Virus expression was assessed using standard histology methods.

For stereotaxic surgeries (Kopf Instruments), a small craniotomy of 0.5 mm in diameter was performed above the injection sites using a microdrill. Injections were performed using a Nanoject III injector and glass capillaries (Süd-Laborbedarf Gaunting); 120nL of virus were injected on each side at a rate of 1nL/sec before optic fibers were slowly inserted and fixed to the skull using Super-Bond (Sun Medical Co. Ltd). Mice were allowed 6 weeks to recover before starting experiments.

### Photometry recordings

Photometry recordings were performed using a camera-based bundle-imaging fiber photometry system (Doric Lenses) using interleaved illumination produced by two LEDs (2 excitation wavelengths: 405 nm and 465 nm / 2 cycles, resolution 1200 ×1200, effective sampling rate 20 Hz). Fluorescence produced by 405-nm excitation provided a real-time control for motion artifacts. Recordings were performed using the Doric Neuroscience Studio software.

### Optogenetics

Optogenetic activation of D1-ChrimsonR and D2-ChrimsonR cells was performed using a red laser (635nm, Laserglow Technologies). Laser output frequency was 4-ms pulses at 20Hz, driven by an Arduino board. Optogenetic inhibition of D1-eNpHR3.0 and D2-eNpHR3.0 cells was achieved with a yellow laser (589 nm, Laserglow Technologies). The laser delivered a constant light when inhibition was triggered and ramped down at the end of an inhibition bout to avoid neuronal activity rebound. D1-GFP and D2-GFP control mice were split evenly between opto-inhibitory and opto-excitatory stimulation patterns.

The lasers were connected to the bilateral fiber implants using a 1×2 fiber-optic rotary joint (Doric Lenses) and yielded an output power of ~8mW at the end of each fiber tip. Laser pulse onset parameters are described in each relevant experiment section.

### ‘Motivation to retrieve a reward’ task

#### Pre-surgery screening

Prior to surgeries, both photometry and optogenetic cohorts mice were screened in this simple reward delivery task. Mice were habituated to the experimental chambers (Mouse Touch Screen Chambers installed in sound-proof chambers, Lafayette Instruments) for 20 min per day for two days. The task schedule was designed and controlled by ABET II software (Lafayette Instrument). On three consecutive days, mice could receive up to 15 milkshake deliveries (Energy milk, strawberry flavor, Emmi Schweiz AG; 20 μL/delivery; to which mice had been previously habituated in the home cage) during a maximum of 30 min. The session ended when whichever of the 15 deliveries or 30 min came first. The inter-trial interval varied pseudo-randomly, 30 ± 10 sec, counted from the moment a mouse exited the food magazine. Only those able to perform the task in 10 min or less were implanted. This was an indication that mice made the association between the cue and food delivery. The house light (white light) was on during the sessions. The tray light was turned on at each milkshake delivery and turned off when the mouse visited the food magazine. Milkshake delivery was associated with a sound cue. Consumption onset was defined as the first food magazine beam break following each delivery.

#### Photometry cohort

The same paradigm used for pre-surgery screening was run again on 3 consecutive days while photometry signals were being recorded (see *Photometry Recordings* section for details).

#### Optogenetics cohort

Here, the task structure was similar to the pre-surgery screening task, except mice performed the task over two days only. The first day was identical to the pre-screening task and served as a baseline. On the second day, the task was identical but 10s of optogenetic stimulation either started from milkshake delivery cue (D1-eNpHR3.0, D2-eNpHR3.0 and GFP controls) or started at food magazine visit onset (D1-ChrismonR, D2-ChrismonR, and GFP controls). After a few-weeks wash-out period this 2-day experiment was repeated, only this time mice which had received optogenetic stimulation during milkshake delivery now received it during the food magazine visit and *vice versa*. In these tasks the activation and inhibition patterns were the same as those described in the *Optogenetics* section, repeated continuously over the 10 sec each stimulation lasted for. Instead of lasers, light was delivered using a PlexBright optogenetic stimulation system (Plexon Inc.) consisting of an Optogenetic Controller (Plexon Inc.) and orange LEDs for activation (compact LED, 620 nm, Plexon Inc.) or lime LEDs for inhibition (compact LED, 550 nm, Plexon Inc.). The output at the end of each patchcord (200/230 μm, 0.5NA, Plexon Inc.) was ~5mW. The patterns were designed in and controlled by Radiant software (Plexon Inc.) which was itself triggered by ABET II software during the task.

### *Ad libitum* consumption experiment

Mice were first habituated to spring-based tip bottles (EBECO) in the home cage using water. Habituation to the arenas (20×20cm^2^ arenas, Habitest modular mouse test cage, Coulbourn Instruments, USA) and patchcords (200 μm, low-autofluorescence, Doric Lenses) was performed on 2 consecutive days before the experiment started. The test cages were placed in closed, sound-attenuating chambers (Coulbourn Instruments, USA) and illuminated using infra-red light only. Licks were detected using a capacitive sensor (Capacitive Touch Breakout, Sparkfun) and recorded at 500Hz throughout the sessions using a National Instruments board (USB-X Series DAQ, NI-6343) and custom built LabView software (National Instruments). Weights consumed were recorded at the end of each session. Mice were not food deprived for these experiments.

#### Photometry cohort

On experimental days, mice were placed in the arenas for 20-min long sessions (one session/day for each mouse) with free access to one spring-based tip bottle (EBECO) containing water (days 1-3), strawberry milkshake (days 4-8). Concomitant photometry signals were recorded as described in the *Photometry Recordings* section above.

#### Optogenetics cohort

Over a 20-min session with access to milkshake, laser stimulation started (according to the patterns described in the *Optogenetics* section) every time a mouse started licking and continued for the duration of licking.

### Hedonic shifting experiment

For this experiment only ChrimsonR mice (and their respective GFP controls) were used. Mice were habituated in the home-cage to two novel flavors of milkshake which were matched for their calorie and macro-nutrient contents: a chocolate-flavored and a vanilla-flavored milkshake (‘Milk Choco Mountain’ Protein, ‘Vanilla Drive’, respectively, Chiefs AG Schweiz).

#### Pre-test phase

this phase served to test the innate preference of mice for one flavor over the other. Testing was performed over two consecutive days with 20-min sessions in the same Coulbourn Instruments chambers as described in the *Ad libitum consumption Experiment* section. Two spring-based tip bottles (EBECO), each with a different flavor of milkshake were freely available. The positions of the bottles in the cage were counterbalanced across mice. Licking and amounts consumed were recorded as described in the *Ad libitum consumption experiment* section. The time spent licking at each bottle on the second day of the pre-test phase was used to determine which was each mouse’s preferred flavour.

#### Conditioning phase

Two 10-min conditioning sessions were run, one in the morning, and one in the afternoon for 7 consecutive days. In each of these sessions only one flavor was available at a time, and the order of the sessions was swapped each day (see **Figure 4a**). Licking at the mouse’s initially-less-preferred flavor triggered optogenetic activation for as long as the mouse kept licking, using the patterns and equipment described in the *Optogenetics* and *Ad libitum consumption experiment* sections respectively.

#### Choice-test phase

24 hours after conditioning, mice were assessed for their preference again. During the choice-test, both flavors were present simultaneously for 20-min with no optogenetic stimulation on either flavor.

Throughout the conditioning and choice-test phases, but not for the initial preference pre-test phase, mice were mildly food-deprived.

### Data Analysis

#### Licking microstructure analysis

Based on previous literature [36, 37], licking behavior was analyzed by looking at the following parameters (see **Figure 1f**):

Single lick durations, defined as an uninterrupted high signal.
Inter-lick intervals (ILI), defined as an uninterrupted low in the signal of up to 250ms. An upper boundary was set in accordance to Johnson [37], to distinguish them from inter-bout intervals.
Number of licking bouts, a bout was defined as a licking episode of at least 500ms, allowing for interruptions within a bout of up to 250ms.
Licking bout duration, length of a given bout in seconds.
Licking bout size, number of individual licks within a given bout.
Inter-bout intervals (IBI), defined as interruptions in licking of more than 250ms allowing for licking occurrences of up to 250ms, i.e. a short lick was not considered a bout and thus does not interrupt a given IBI.

For the regression analysis (see below), licking bouts had a slightly more inclusive definition, starting with at least 100 ms of licking and allowing for pauses of up to 1 sec.

#### Multiple regression analysis

Correlations between photometry signals and ‘liking’/‘wanting’ parameters were performed using the SPSS (IBM Corp. Version 28) linear regression tool. A subset of 8 parameters depicting the photometry signal around a licking bout onset was selected: maximum, minimum and area under the curve (AUC) in the 5 secs following lick onset; mean and median signal over the 3 sec before lick onset; mean and median signal over the 3 sec following lick onset and standard deviation of the non-z-scored signal in the 5 sec preceding lick onset. Because aiming to fit too many variables in a regression model can lead to spurious results, we used all parameters simultaneously in a first step only, to assess correlations between predictors and see which predictors had the highest and most significant correlations with the outcome variable. If predictors were highly correlated between themselves (Pearson’s r >8.5, VIF>10, tolerance<0.1, checking expression of each predictor in eigen vector base) then only the predictor with the most significant correlation with the outcome variable was kept. Then, in a second step, another linear regression was performed keeping only the non-collinear predictors. A final block-wise (hierarchical) regression was done keeping only the predictors that showed significant correlations with the outcome variable and/or model coefficients significantly different from 0. This final block-wise approach – which started with the predictor yielding the highest significant standardized model parameter (β) and continued in decreasing order of βs – allowed us to evaluate from which predictor adding one more predictor to the model did not yield better predictive power (looking at the p value associated to the F-ratio change). Predictors whose addition to the model gave a significant F-ratio change (i.e., improved the model) are reported in **Figures S2-4**. Case-wise diagnostics allowed us to evaluate the adequacy of our dataset, namely whether 95% of data points had standardized residuals within ±2, and around 1% of the sample at most had standardized residuals over 2.5. Assumptions of the model (absence of heteroscedasticity, linearity) were evaluated by plotting standardized residuals against standardized predicted values. Normal distribution was evaluated via the observed vs. predicted cumulative probability plots. The Durbin-Watson index was used to evaluate whether the model’s residuals were independent (index should be as close to 2 as possible, and at least >1 and <3). As our aim here was to evaluate which aspects of the photometry signal (i.e., neuronal activity) best predicted ‘liking’ and/or ‘wanting’, but not to apply the model to future datasets, we did not use bootstrapping. Nonetheless, to assess the validity of our model for a general population, cross-validation was performed using the ‘adjusted R^2^ value.’ A value as close as possible to R^2^ indicated that the model generalizes well to the population beyond the sample data.

#### ‘Motivation to retrieve a reward’ task analysis

When analyzing photometry signals around food delivery, trials where the mouse entered the food magazine within 5 sec after food delivery cue were not included so that signals did not include food consumption, but only food anticipation (**Figure 3d**). For analyses of signals around food magazine visits, trials for which the mouse exited the food magazine just after entering it were discarded (**Figure 3e**). When calculating the average food magazine visit length, visits of less than 2 seconds were excluded to avoid skewing the mean to short values of very brief – and mostly non-consummatory – visits (**Figure 3j,k**).

#### Statistical analysis

Data were analyzed using Matlab R2019b (MathWorks). All statistical tests and descriptive statistics were performed using SPSS software (IBM Corp. Version 28). Sex was taken as a factor in the analysis but did not reveal significant differences in all comparisons. Therefore, since sex was not the primary independent variable of interest in this study, and given the lack of statistical differences, we pooled data from both sexes for all further analyses. Tests and their results are presented in the figure legends. Unless otherwise stated, data are presented as mean ± SEM, and a p-value ≤ 0.05 was considered to indicate significance. In the motivation task (**Figure 3f-k**), outlier trials (defined as points whose absolute z-score is >3.29) were excluded from the statistical analysis.

## Author Contributions

Conceptualization, D.P.R.; Investigation, M.C.C.G.; Formal Analysis, M.C.C.G., P.V. and D.P.R.; Writing – Original Draft, M.C.C.G. and D.P.R.; Writing – Review & Editing, M.C.C.G., E.B., D.B. and D.P.R.; Visualization, M.C.C.G.; Methodology, M.C.C.G. and D.P.R.; Funding acquisition, D.B. and D.P.R.; Supervision, D.P.R.

## Acknowledgments

We would like to thank Valeria Franco Betzler for her assistance in histological evaluations. This study was supported by the Swiss National Science Foundation (grant no. 310030_189110 to DPR and DB) and the Novartis Foundation for medical-biological Research (grant no. 21A061 to DPR).

## Supplementary figures

**Figure S1.**
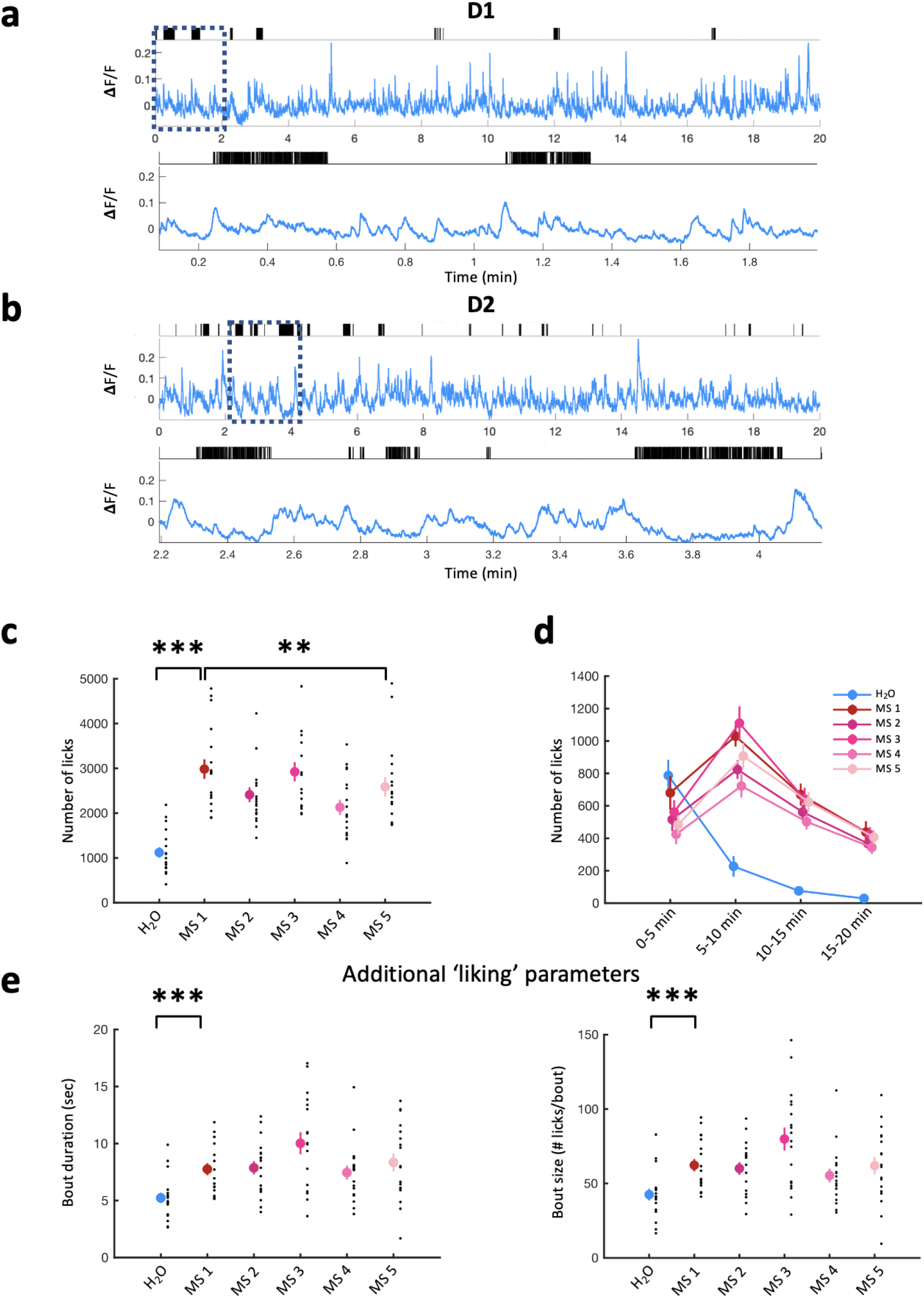
An increase in palatability is reflected in hedonic behavior (‘liking’) parameters. (a) Example photometry recording from a 20-min session with access to water from a D1-GCaMP6s mouse. Licks indicated by black raster plot above. Bottom, zoom of photometry signal and licks showing only 2 min indicated by black dotted square above. Photometry signals are (465nm signal – fitted 405 signal)/fitted 405nm signal. (b) As in (a) for a D2-GCaMP6s mouse. (c) Lick number during daily 20-min sessions with access to water (H_2_O, first session, blue dot) and milkshake (MS - 5 sessions, red and pink dots). Mean ± SEM values are plotted in color, and individual values from each mouse in black. D1-GCaMP6s and D2-GCaMP6s mice are pooled, n=18. One-way repeated measures ANOVA: water vs. MS 1 (F(1,17)=147.327, p<0.001), MS 1 to MS.5 (F(4,68)=19.895, p<0.001), MS 1 vs. MS 5 (F(1,17)=11.006, p=0.004). (d) Lick number over 5-min blocks across daily 20-min sessions with access to water (first session) and MS (5 sessions). Values are plotted (in color) as mean ± SEM, n=18. (e) Additional parameters calculated from lick recordings canonically associated with ‘liking’: bout duration (left) and bout size (i.e., number of licks per bout, right) across one session with access to water and 5 sessions with access to MS. D1-GCaMP6s and D2-GCaMP6s mice are pooled. Mean ± SEM values are plotted in color, and individual values from each mouse in black, n=18. One-way repeated measures ANOVA, Bout duration: water vs. MS 1 (F(1,17)=25.529, p<0.001), MS 1 to MS 5 (F(4,68)=4.245, p=0.004), MS 1 vs. MS 5 (F(1,17)=0.675, p=0.423); Bout size: water vs. MS 1 (F(1,17)=26.825, p<0.001), MS 1 to MS 5 (F(4,68)=5.282, p=0.001), MS 1 vs. MS 5 (F(1,17)=0.003, p=0.956) MS: Milkshake. P-values reported on the figures as follows: **p≤0.01, ***p<0.001.

**Figure S2.**
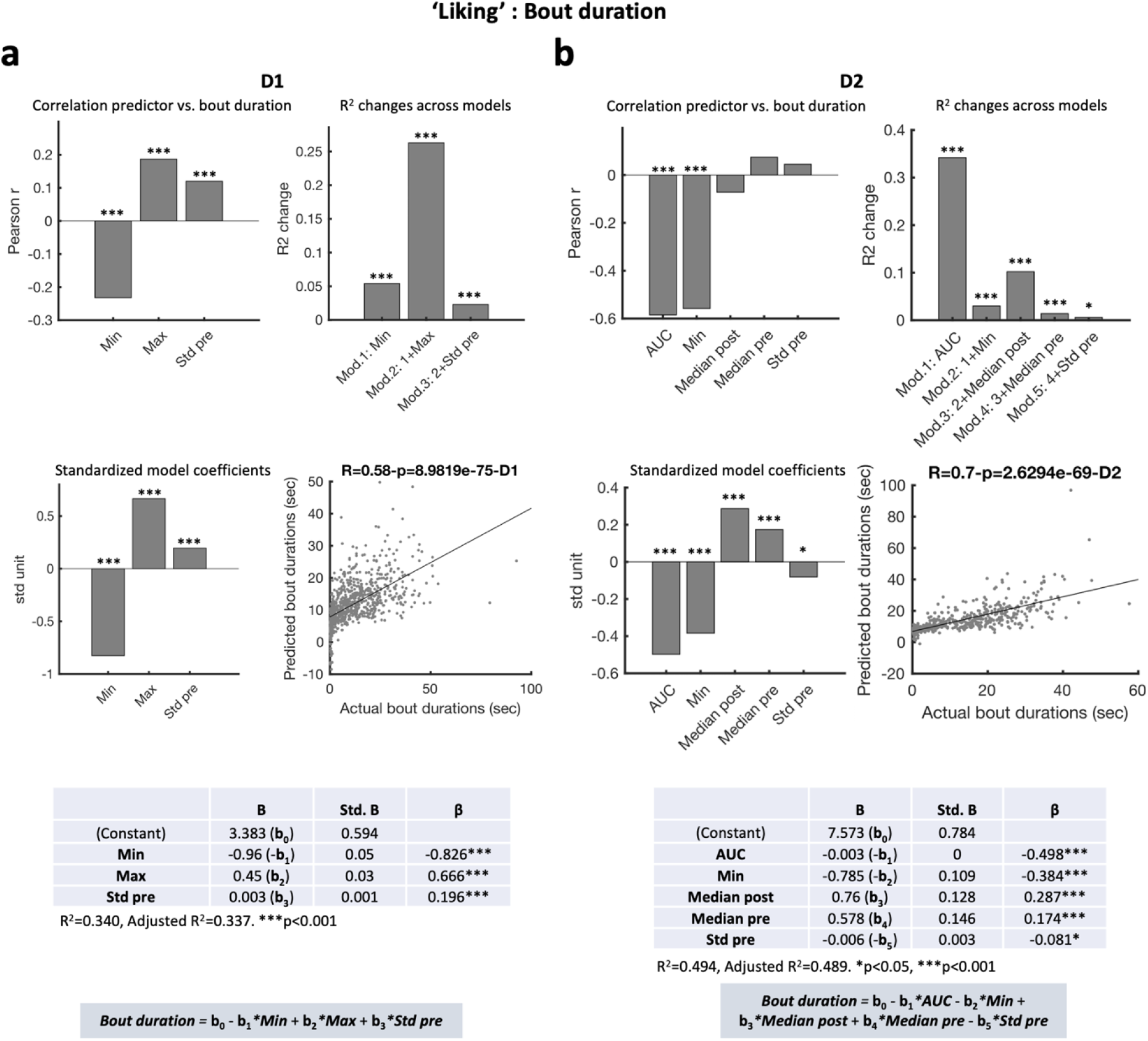
Photometry signal features of NAc D1 and D2 cells around *ad libitum* palatable food consumption are good predictors of ‘liking’ of the food consumed. (a)Multiple regression analysis using variables extracted from photometry signals for D1-GCaMP6s mice (n=6) around licking onset to predict licking bout duration. Top left: Pearson’s r values of single correlations between predictors and bout duration (Min: Minimum value in the 5 sec following lick onset, Max: Maximum value in the 5 sec following lick onset, Std: standard deviation of the non-z-scored signal in the 5 sec preceding lick onset – see Methods). Top right: R^2^ change evaluating whether a given regression model (Mod.) significantly improves the prediction of bout duration compared to the previous model; the p-value associated with the R^2^ change is reported above each bar. The R^2^ change indicates whether the amount of variance in the outcome (bout duration) can be explained by the added predictor. For model 1, Min was the only predictor included. The R^2^ change shows that using the Min to predict bout duration significantly improves predictions compared to using the grand mean of bout duration. For model 2, Min and Max were included as predictors. The R^2^ change shows that including the Max of the photometry signal around licking onset in addition to the Min significantly improves the model’s performance in predicting bout duration, with similar reasoning for subsequent models (Model 3). Middle left: Standardized coefficients of the final model; standardized coefficients allow to compare the relative contribution of each predictor, regardless of the units used to measure them. The higher the absolute value of a standardized coefficient, the higher the contribution to the model. The p-values associated with these contributions (i.e., whether they are significant, regardless of their magnitude) is reported above each bar. Middle right: Plot of the actual vs. predicted bout duration with the R and p values reported on top. Bottom: model coefficients (B), standard deviation for the coefficients (Std. B) and standardized model coefficient (β) of each predictor in the final model, of which the equation is reported below the table. (b) As in (a) but for D2-GCaMP6s mice (n=5). AUC: area under the curve over the 5 sec following lick onset, Median pre: median signal over the 3 sec before lick onset, Median post: median signal over the 3 sec following lick onset. AUC: area under the curve, Max: maximum, Min: minimum, pre: before licking onset, post: following licking onset, Std: standard deviation. p-values reported on the figures as follows: *p≤0.05, ***p<0.001.

**Figure S3.**
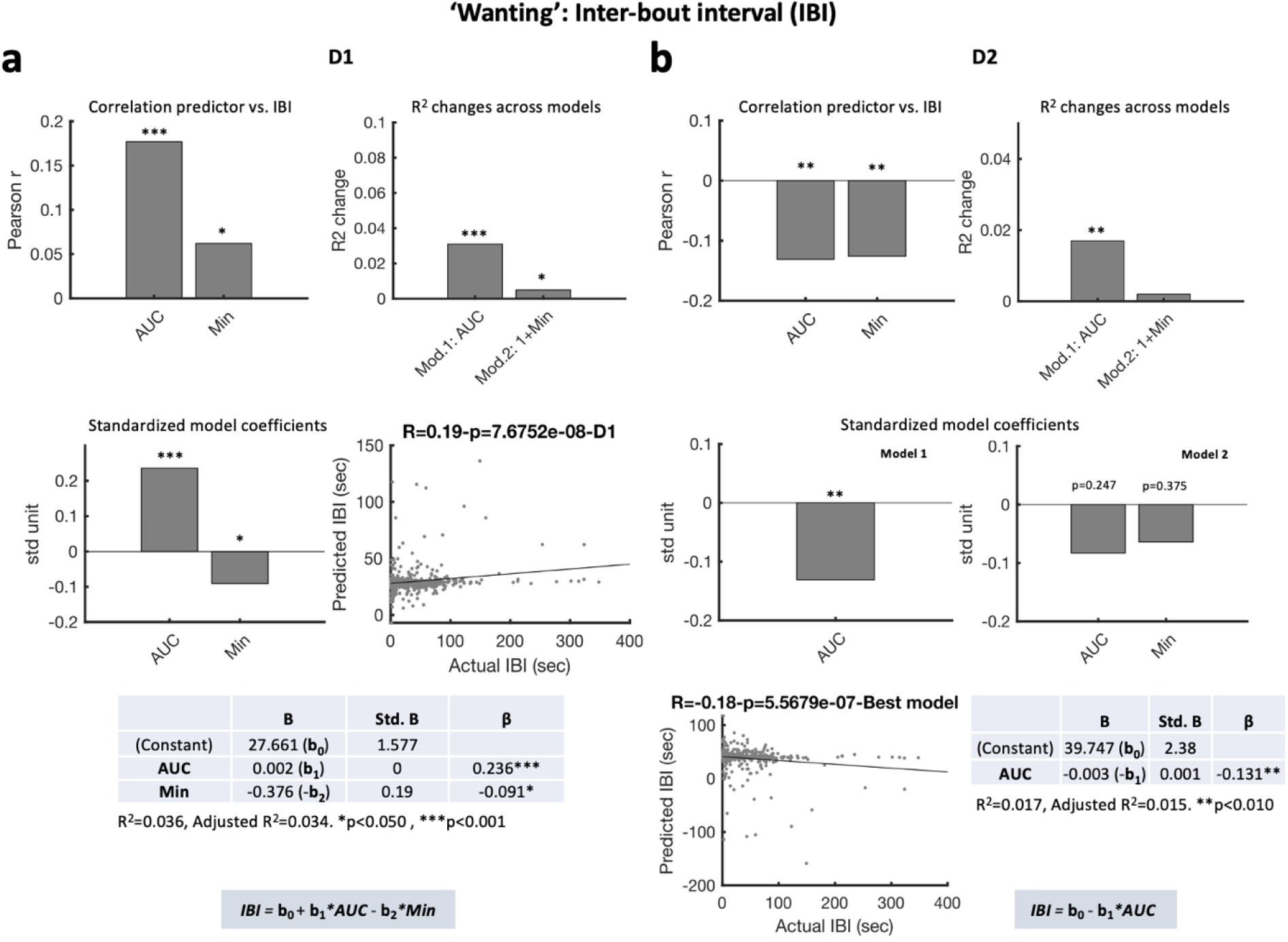
Photometry signal features of NAc D1 and D2 cells around *ad libitum* palatable food consumption are not good predictors of ‘wanting’ of the food consumed. (a) As in Figure S2a but for prediction of inter-bout intervals (IBI) in D1-GCaMP6s mice (n=6). (b) As in (a) but for D2-GCaMP6s mice (n=5). Note that here although both the AUC and Min predictors were individually significantly correlated with IBI (top left), only the model considering the AUC yielded a significant improvement in predicting IBI duration compared to using the grand mean of IBI duration (i.e., no better predicting power of model 2 compared to model 1) (top right and middle right). AUC: area under the curve, IBI: inter-bout interval, Min: minimum, Std: standard deviation. p-values reported on the figures as follows: *p≤0.05, **p≤0.01, ***p<0.001.

**Figure S4.**
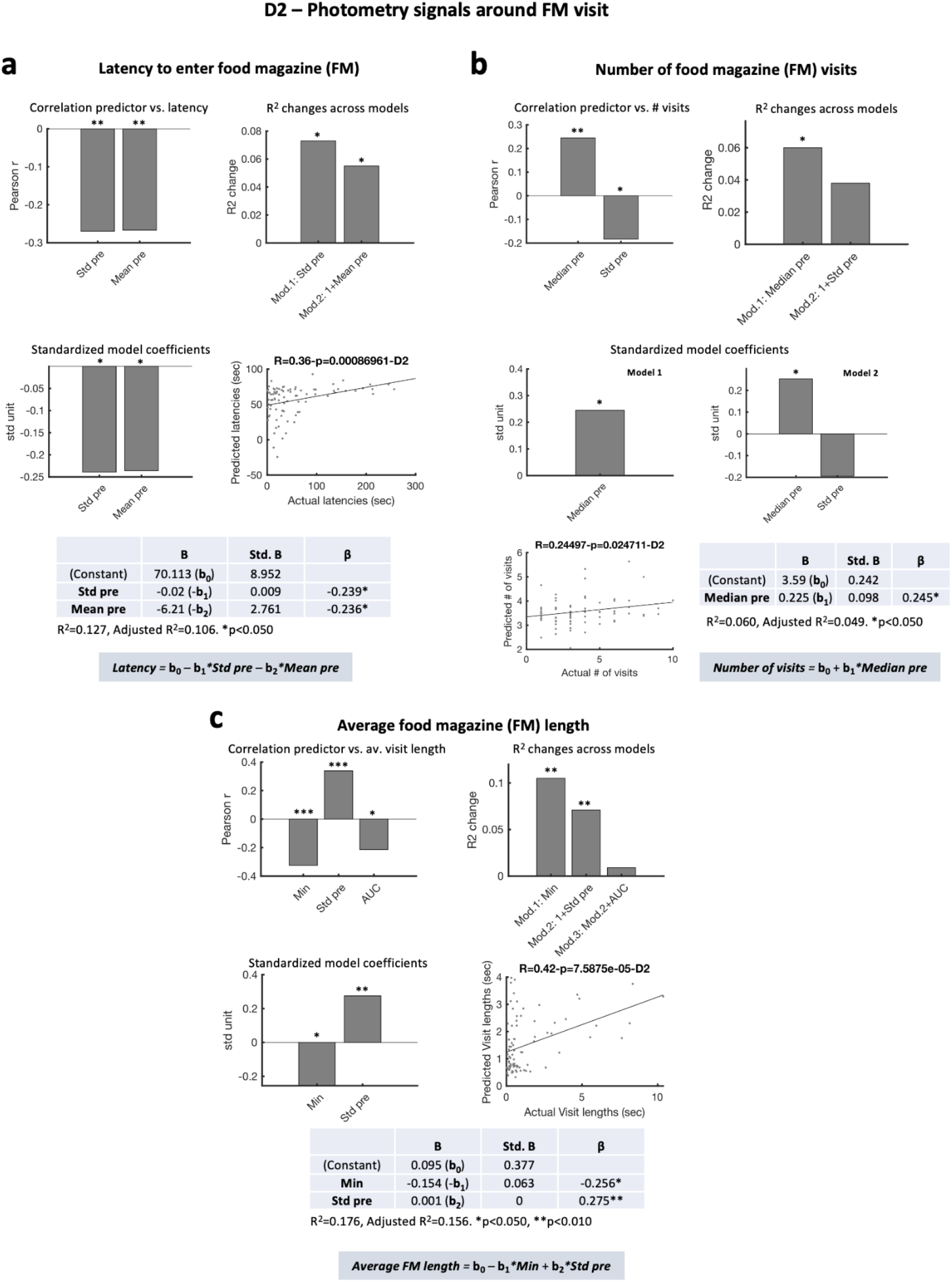
The decrease in D2 cell activity during consumption favors longer licking bouts. (a) Multiple regression analysis using variables extracted from the photometry signals for D2-GCaMP6s mice around food magazine (FM) visit onsets to predict the latency to enter the FM following a food delivery. Top left: Pearson’s r values of single correlations between predictors and FM visit latency (Mean pre: mean signal over the 3 sec before FM visit onset, Std pre: standard deviation of the non-z-scored signal over the 5 sec preceding FM visit onset – see Methods). Top right: R^2^ change evaluating whether a given regression model (Mod.) significantly improves the prediction of FM visit latency compared to the previous model; the p-value associated with the R^2^ change is reported above each bar. The R^2^ change indicates whether the amount of variance in the outcome (latency to enter the FM) can be explained by the added predictor. For model 1, Std pre was the only predictor included. The R^2^ change shows that using the Std pre to predict latency significantly improves predictions compared to using the grand mean of latencies. For model 2, Std pre and Mean pre were included as predictors. The R^2^ change shows that including the mean photometry signal over the 3 sec before FM visit in addition to the Std pre significantly improves the model’s performance in predicting latency to enter the FM. Middle left: Standardized coefficients of the final model; standardized coefficients allow to compare the relative contribution of each predictor, regardless of the units used to measure them. The higher the absolute value of a standardized coefficient, the higher the contribution to the model. The p-values associated with these contributions (i.e., whether they are significant, regardless of their magnitude) is reported above each bar. Middle right: Plot of the actual vs. predicted FM visit latency with the R and p values reported on top. Bottom: model coefficients (B), standard deviation of the coefficients (Std. B) and standardized model coefficient (β) of each predictor in the final model, of which the equation is reported below the table. (b) As in (a) but for prediction of number of food magazine visits. Median pre: median signal over the 3 sec before FM visit onset. Note that here although both the Median pre and Std pre predictors were individually significantly correlated with the number of FM visits (top left), only the model considering the Median pre yielded a significant improvement in predicting number of FM visits compared to using the grand mean of the number of visits (i.e., no better predicting power of model 2 compared to model 1) (top right and middle right). (c) As in (a) but for the average food magazine visit length. Min: Minimum value in the 5 sec following FM visit onset. AUC: area under the curve. AUC: area under the curve. FM: food magazine, Min: minimum, pre: before FM visit onset, Std: standard deviation. p-values reported on the figures as follows: *p≤0.05, **p≤0.01, ***p<0.001.

**Figure S5.**
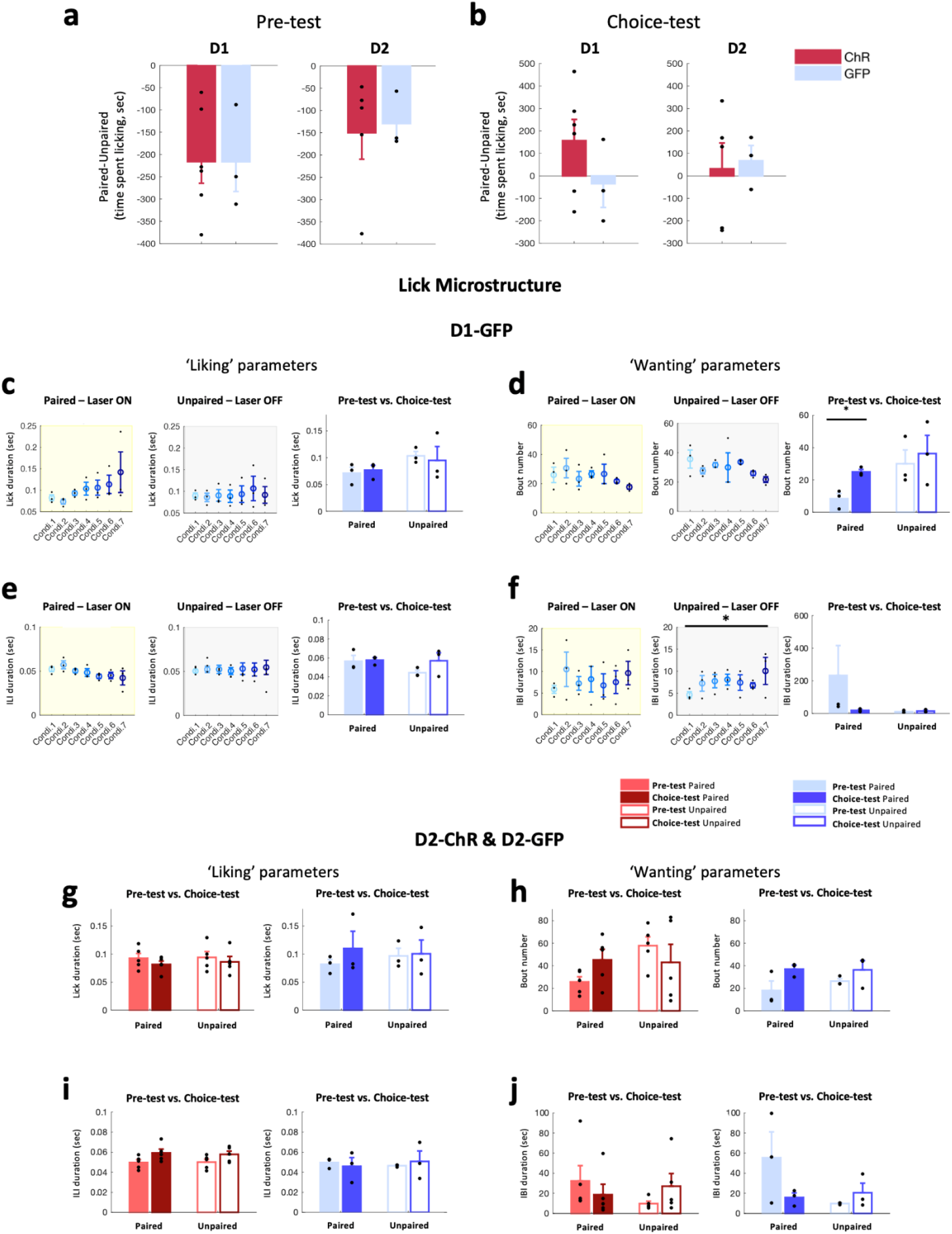
Activation of D2 cells does not lead to a shift in flavor preference for the optoactivation-paired flavor. (a) Difference in the time spent licking the paired and unpaired flavors during the pre-test session of the hedonic shifting experiment in D1-cre (left) and D2(A2a)-cre mice (right) expressing ChrimsonR (ChR, red) or GFP (blue). One-way ANOVA (opsin), D1: F(1,7)<0.001, p=0.994, D2: F(1,6)=0.062, p=0.812. (b) As in (a) but for the time spent licking each flavor on the choice-test session. One-way ANOVA (opsin), D1: F(1,7)=1.526, p=0.257, D2: F(1,6)=0.048, p=0.833. (c) Single-lick duration calculated from lick recordings indicating ‘liking’ across the conditioning (Condi.) sessions (left & middle panels) and pre-test & choice-test sessions (right panel) in the paired and unpaired flavors in D1-GFP mice. Mean ± SEM values are plotted in color, and individual values from each mouse in black. RM-ANOVA one-sided: Conditioning, paired flavor (day: F(6,12)=1.705, N.S.), unpaired flavor (day: F(6,12)=0.405, N.S.); Choice-test vs. pre-test, paired flavor (test day: F(1,2)=3.865, N.S.), unpaired flavor (test day: F(1,2)=0.224, N.S.). (d) As in (c) but for the number of licking bouts. RM-ANOVA – one-sided: Conditioning, paired flavor (day: F(6,12)=0.815, N.S.), unpaired flavor (day: F(6,12)=0.974, N.S.); Choice-test vs. pre-test, paired flavor (test day: F(1,2)=12.563, p=0.036), unpaired flavor (test day: F(1,2)=0.113, N.S.). (e) As in (c) but for inter-lick interval (ILI) durations. RM-ANOVA – one-sided: Conditioning, paired flavor (day: F(6,12)=1.528, N.S.), unpaired flavor (day: F(6,12)=1.128, N.S.); Choice-test vs. pre-test, paired flavor (test day: F(1,2)=0.044, N.S.), unpaired flavor (test day: F(1,2)=3.277, N.S.). (f) As in (c) but for the inter-bout interval (IBI) durations. RM-ANOVA – one-sided: Conditioning, paired flavor (day: F(6,12)=0.689, N.S.), unpaired flavor (day: F(6,12)=13.824, p=0.033); Choice-test vs. pre-test, paired flavor (test day: F(1,2)=1.307, N.S.), unpaired flavor (test day: F(1,2)=0.183, N.S.). (g) Single-lick duration during the pre-test & choice-test sessions in the paired (full bars) and unpaired (empty bars) flavors in D2-ChR (left, red) and D2-GFP (right, blue) mice. Mean ± SEM values are plotted in color, and individual values from each mouse in black. RM-ANOVA: Choice-test vs. pre-test, D2-ChR: paired flavor (test day: F(1,4)=1.307, N.S.), unpaired flavor (test day: F(1,4)=0.599, N.S.); D2-GFP: paired flavor (test day: F(1,2)=0.517, N.S.), unpaired flavor (test day: F(1,2)=0.011, N.S.). (h) As in (g) but for the number of licking bouts. RM-ANOVA: Choice-test vs. pre-test, D2-ChR: paired flavor (test day: F(1,4)=2.538, N.S.), unpaired flavor (test day: F(1,4)=0.423, N.S.); D2-GFP: paired flavor (test day: F(1,2)=7.367, N.S.), unpaired flavor (test day: F(1,2)=0.906, N.S.). (i) As in (g) but for inter-lick interval (ILI) durations. RM-ANOVA: Choice-test vs. pre-test, D2-ChR: paired flavor (test day: F(1,4)=6.971, N.S.), unpaired flavor (test day: F(1,4)=1.887, N.S.); D2-GFP: paired flavor (test day: F(1,2)=0.141, N.S.), unpaired flavor (test day: F(1,2)=0.153, N.S.). (j)As in (h) but for the inter-bout interval (IBI) durations. RM-ANOVA: Choice-test vs. pre-test, D2-ChR: paired flavor (test day: F(1,4)=0.394, N.S.), unpaired flavor (test day: F(1,4)=2.593, N.S.); D2-GFP: paired flavor (test day: F(1,2)=2.962, N.S.), unpaired flavor (test day: F(1,2)=1.342, N.S.). ChR: ChrimsonR, Condi.: conditioning, IBI: inter-bout interval, ILI: inter-lick interval. p-values reported on the figures as follows: *p≤0.05.

